# Tuning viscoelasticity of dynamic covalent hydrogels for human tissue modeling

**DOI:** 10.1101/2025.10.16.682916

**Authors:** Narelli de Paiva Narciso, Fotis Christakopoulos, Michelle S. Huang, Neil J. Baugh, Carla Huerta-López, Eliana X. Matos, Kristine P. Pashin, Andrew Spakowitz, Sarah C. Heilshorn

**Author notes:** **Correspondence** Corresponding author Sarah C. Heilshorn.

## Abstract

The development of three-dimensional (3D) *in vitro* tissue culture models is critical for biomedical research. Hydrogel-based systems have become a preferred scaffold for 3D models, as they have tunable viscoelastic properties, which are well-known to influence cell morphology and function. In particular, reversible hydrogel crosslinks formed through dynamic covalent chemistry (DCC) can introduce viscoelastic behavior including stress relaxation. However, traditional strategies to increase stress-relaxation rates in DCC gels rely on faster bond kinetics, resulting in faster erosion rates that prevent their use for long-term 3D culture. As an alternative strategy, we explore the use of molecular parameters (specifically molecular weight and degree of functionalization) to independently control the stiffness and stress relaxation behavior while preventing rapid erosion. As demonstration, we develop and validate a modified theoretical model of gel viscoelasticity applied to a two-component DCC gel composed of modified hyaluronic acid and elastin-like protein. Finally, we utilize this tunable gel platform to explore the impact of scaffold viscoelasticity on encapsulated human neural progenitor cells. In summary, this work expands the molecular design space of DCC hydrogels to achieve tunable viscoelastic properties for 3D *in vitro* models.

## 1 INTRODUCTION

The development of *in vitro* tissue culture models has been paramount in biomedical research, as they allow for carefully controlled variables, mechanistic studies, potential for high-throughput screening, and a decrease in our reliance on animal models.Tibbitt and Anseth (2009), Xu et al. (2022), Terzopoulou et al. (2022), Leung et al. (2022) Thus, there is a growing interest in so-called “new approach methodologies” (*i.e*. NAMs) such as organ-on-a-chip systems, 3D cell culture models, and patient-derived organoids that can improve our understanding of human development, disease progression, drug discovery, and regenerative medicine Leung et al. (2022), Huang et al. (2024), Ingber (2022), Huang et al. (2025a), Kim et al. (2020), Tang et al. (2022), Liu et al. (2025a), Abuwatfa et al. (2024). However, a central challenge in the field is the ability to reproducibly emulate the complex biomechanical and *Journal* 2025;00:1–18 biochemical environment of native tissue matrices.Elosegui-Artola et al. (2023), Tibbitt and Anseth (2009), Abuwatfa et al. (2024)

One of the key challenges is mimicking the viscoelastic mechanical properties of natural tissue, which present a time-dependent response to stress and deformation that can be exerted either by cells or by external stimuli Chaudhuri et al. (2015), Courbot and Elosegui-Artola (2025), Huang et al. (2019), Elosegui-Artola et al. (2023), Dai et al. (2025). The importance of mechanical cues to guide cell behavior have been widely reported, impacting stem cell differentiation, cell migration, morphology, and proliferation, among other processes.Cheng et al. (2022), Hachet et al. (2012), Hunt et al. (2021), LeSavage et al. (2024), Navarro et al. (2022), Darnell et al. (2017), Dai et al. (2025) More recently, the importance of the kinetics of stress-relaxation has been highlighted, in addition to the more traditionally studied stiffness (storage modulus, *G′*) Chaudhuri et al. (2015 2016 2020), Elosegui-Artola et al. (2023), Darnell et al. (2017). Across different tissues and stages of development or disease progression, both stiffness and stress-relaxation rate can significantly change to impact cell behavior. Chaudhuri et al. (2015). As a consequence, hydrogels have risen to prominence as ideal *in vitro* systems for studies of cellular biomechanics due to their tunable mechanical properties (Tibbitt and Anseth (2009), Xu et al. (2022)). While the effects of hydrogel stiffness on cellular behavior have been extensively explored,Cheng et al. (2022), Hachet et al. (2012), Hunt et al. (2021), LeSavage et al. (2024), Navarro et al. (2022), Liu et al. (2025b) studies on the consequences of different stress relaxation rates, especially in combination with varied stiffness, are less common. A key reason for this is the inherent difficulty of independently tuning stiffness and stress-relaxation kinetics within a cell-compatible hydrogel. Moreover, stress-relaxing hydrogels tend to suffer from rapid erosion profiles (often on the order of 1-3 days), preventing longer cell culture times. Hull et al. (2023), Borelli et al. (2022)

Our group has previously utilized recombinant biopolymers and dynamic covalent chemistry (DCC) to achieve hydrogels with tunable stiffness, stress relaxation rate, injectability, and cell-instructive domains.de Paiva Narciso et al. (2023), Roth et al. (2023), Huang et al. (2025b), Suhar et al. (2023). These materials are formed through a recombinant elastin-like protein (ELP) and recombinant hyaluronic acid (HA), which together we term HELP gels. Recombinant biopolymers offer the advantages of being biodegradable and cell-interactive (similar to native biopolymers) while also being reproducible and produced with animal-free methods (similar to synthetic polymers).de Paiva Narciso et al. (2023), Hefferon et al. (2023), Wang et al. (2018), LeSavage et al. (2018) The ELP protein incorporates alternating blocks of repetitive elastin-like peptides that provide mechanical elasticity and cell-interactive sequences derived from fibronectin that provide adhesion to integrin cell-surface receptors Hefferon et al. (2023), Suhar et al. (2023).

HA is one of the main components of ECM throughout the human body and can be chemically functionalized through bioorthogonal chemistry.Xu et al. (2012), Hachet et al. (2012), Madl and Heilshorn (2018) The use of recombinant HA of specific molecular weights allows us to explore the role of molecular parameters in dictating the macroscale hydrogel mechanics.

A commonly reported method to achieve DCC hydrogels with faster stress-relaxation rates is to select crosslinks with faster on-and-off rate kinetics.Hull et al. (2023), Zhang et al. (2023), Hafeez et al. (2018), Han et al. (2022), Anseth and Klok (2016), Muir and Burdick (2021) While successful, these strategies also affect other hydrogel characteristics that are crucial for *in vitro* modeling, such as gelation timeLiu et al. (2024) and erosion kineticsHull et al. (2023). When gelation time is too rapid or too slow, this can prevent homogeneous cell encapsulation; and when erosion is too rapid, this can prevent long-term cell culture. As an alternative approach, we hypothesized that molecular manipulation of the polymer molecular weight and/or degree of functionalization would enable formulation of DCC hydrogels with independent control of stiffness and stress relaxation rate. To test this hypothesis, we synthesized a large family of recombinant HELP hydrogels crosslinked through hydrazone DCC. Through experimental and computational rheological analysis, we identify the molecular design parameters and polymer physics principles that enable tuning and independent control of the hydrogel stiffness and stress-relaxation rate without impacting erosion rate. To demonstrate the potential use of these materials for *in vitro* tissue modeling, we explore how viscoelastic network properties impact the morphology of human neural progenitor cells (NPCs) over 7 days of culture. Given the location of endogenous NPCs in the dentate gyrus of the brain, *in vitro* models are critical to further study of human brain development and disease progression Stiles and Jernigan (2010), Liu et al. (2023).

## 2 RESULTS AND DISCUSSION

### 2.1 Tuning stress relaxation through reaction kinetics

The hydrogels used in this work are comprised of recombinant HA and ELP, termed HELP hydrogels, with the total polymer weight percentage being kept constant at 2 wt% (1 wt% HA and 1 wt% ELP) in all formulations. We chose a recombinant ELP variant that incorporates the fibronectin-derived RGD amino acid sequence that provides cell-adhesive interactions (Figure 1A). LeSavage et al. (2018), Suhar et al. (2023), Hefferon et al. (2023) To enable dynamic crosslinking, the ELP is chemically modified with hydrazine (HYD) moieties, as described in our previous work. Hefferon et al. (2023), de Paiva Narciso et al. (2023) Similarly, HA was functionalized with either aldehyde (ALD) (Figure 1B) or benzaldehyde (BZA) (Figure 1C), to form dynamic hydrazone bonds with the HYD displayed by ELP (Figure 1D). The reaction kinetics for the ALD-HYD are known to be faster than that for the BZA-HYD, resulting in different rates of dynamic bond formation and dissociation.Liu et al. (2024), McKinnon et al. (2014) To evaluate the effect of the reaction kinetics on the bulk hydrogel stiffness, we used three formulations with different ratios of HA-ALD and HA-BZA (100:0, 50:50, 0:100), while keeping the HA *M*_*w*_ constant (60 kDa). For HELP gels formulated with 6 % functionalized HA, all gels had a plateau storage modulus (*G′*) of about 200 Pa, independent of the ALD:BZA ratio (Figure 1E). We then formulated a family of stiffer gels by increasing the HA functionalization to 12 %. As before, we observed that the ALD:BZA ratio had a small but negligible impact on final gel stiffness, with *G′* of about 900 Pa.

**FIGURE 1:**
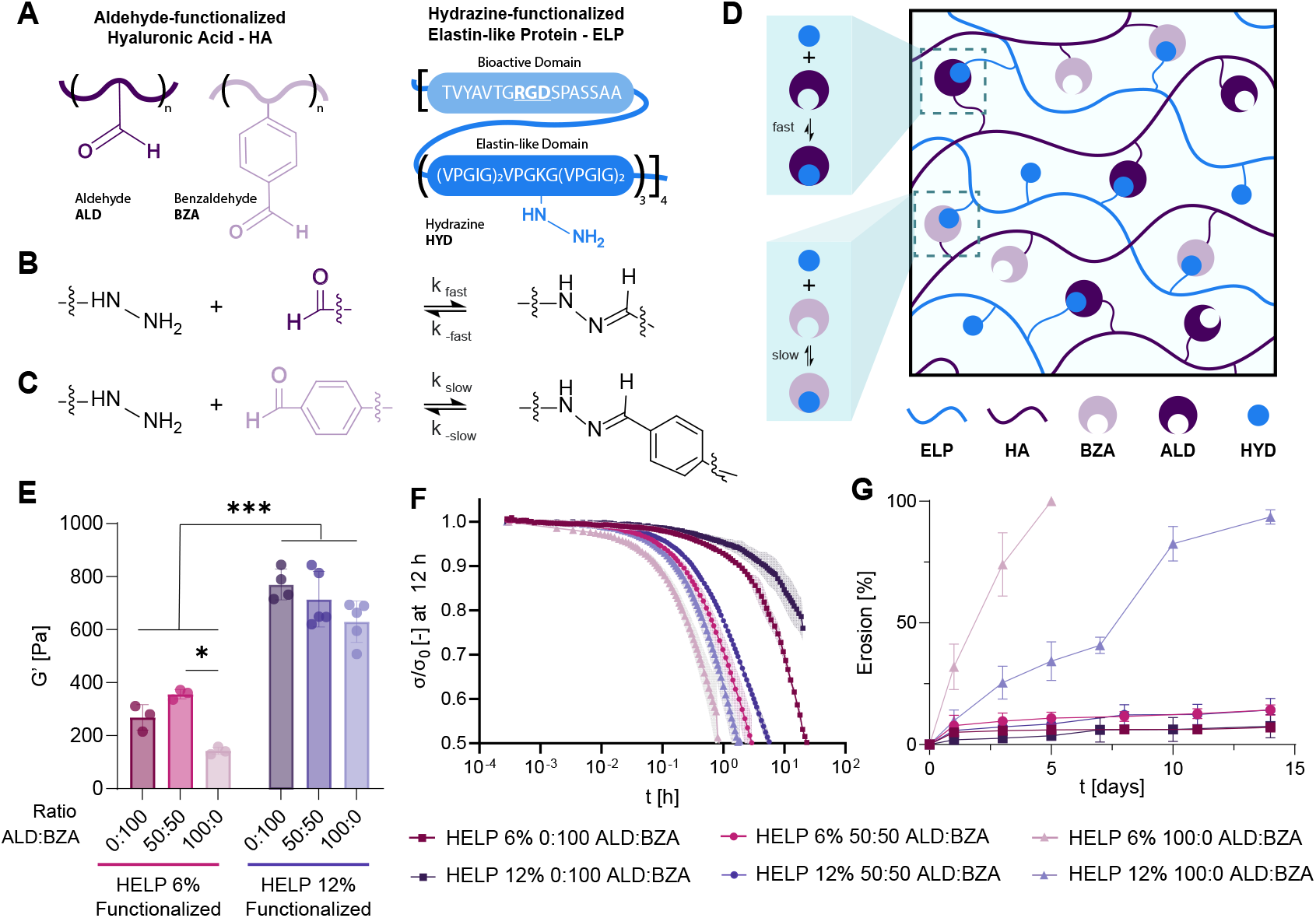
Reaction kinetics control stress-relaxation kinetics and erosion profile. **A**. Schematic of hyaluronic acid (HA) functionalized with aldehyde (ALD) or benzaldehyde (BZA), and the elastin-like protein (ELP), including amino acid sequence, functionalized with hydrazine (HYD). **B**,**C** Representation of ELP-HYD and HA-ALD (B) and HA-BZA (C) forming a dynamic hydrazone bond with fast and slow reaction kinetics, respectively. **D**. Schematic of hydrogel network formed upon mixing HA-ALD and HA-BZA with ELP-HYD, termed HELP gel. **E**,**F**. Storage modulus (*G′*) at an angular frequency of 1 rad s^−1^ (**E**) and stress-relaxation kinetics (**F**) of HELP hydrogels with different ratios of HA-ALD:HA-BZA, consisting of HA with a *M*_*w*_ of 60 kDa and a total amount of functionalization of either 12 % (purple) or 6 % (green). **G**. Erosion profile of HELP gels consisting of 60 kDa HA and with a total functionalization of 6 % and 12 % with either ALD (triangles) or BZA (squares). **E**. One-way ANOVA, p<0.05 *, p<0.01 **, p<0.001 ***, p<0.0001 ****. Specific p-values are listed on Table S1.

In contrast, the stress-relaxation kinetics were found to strongly depend on the bond kinetics (Figure 1F). The higher the ALD:BZA ratio, the faster the stress-relaxation kinetics, with the gels having only the HA-ALD variant (100:0) dissipating 50 % of the applied stress after about 2000 s, and 4000 s, for the 6 % and 12 % functionalized HA, respectively. In contrast, when only BZA-functionalized HA is used (0:100), the relaxation kinetics are considerably slower; even after 12 hours, the 12 % functionalized HA HELP gel had only dissipated about 20 % of the applied stress.

As expected, the differences in ALD:BZA reaction kinetics not only affected the time-scale of stress-relaxation, but also the rate of hydrogel erosion (Figure 1G). HELP hydrogels comprising only HA-ALD (100:0, 12 % functionalization) erode considerably over three days (25 %) and almost completely after 14 days, while those consisting of HA-BZA (0:100, 12 % functionalization) retain about 90 % of their initial mass over the same time-frame. For HELP gels with the lower 6 % functionalization, the erosion rate was even more rapid. Similar rapid erosion rates have been reported for other DCC-crosslinked hydrogels.Borelli et al. (2022), Lin et al. (2024), Anseth and Klok (2016), Jaeschke et al. (2024) This rapid erosion hinders the utility of these materials as scaffolds for *in vitro* tissue models. Thus, while tuning of the crosslinker reaction kinetics is a straightforward way to achieve faster stress-relaxing hydrogels, these materials have practical limitations due to the interplay between stress-relaxation and erosion kinetics.

### 2.2 Impact of molecular weight on stiffness and stress-relaxation kinetics

As an alternative to the use of crosslinking reaction kinetics to control the HELP gel stress relaxation rate, we next explored how HA *M*_*w*_ could alter HELP gel mechanics while keeping the ALD:BZA ratio and degree of functionalization constant (0:100 and 12 %, respectively) (Figure 2A). We observed that the HA *M*_*w*_ did have an appreciable impact on both hydrogel stress relaxation rate (Figure 2B) and stiffness (Figure 2C). Consistent with previous results, increasing the *M*_*w*_ results in an increase in hydrogel stiffness (*G′*) due to an increase in polymer entanglements (Figure 2C).de Paiva Narciso et al. (2023), Cai et al. (2022) Interestingly, an inverse parabolic relationship was observed when evaluating the effect of HA *M*_*w*_ on network stress-relaxation kinetics (Figure 2D-E). Increasing HA *M*_*w*_ from 20 to 60 kDa resulted in slower stress relaxation kinetics (Figure 2B,D-E). This decrease is expected when the relaxation time (*τ*) is governed by entanglements between neighboring polymers, similar to that observed for physical networks and polymer melts, where relaxation time scales as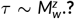Rosa and Winter (1994), Oelschlaeger et al. (2013) However, as HA *M*_*w*_ was further increased to 100 kDa and higher, the stress-relaxation kinetics began to speed up again. This phenomenon is highlighted by observing that the stress relaxation profiles of the 20-kDa and 500-kDa HELP gels are similar (Figure 2B) despite the 500-kDa HELP gel being x-fold stiffer (Figure 2C). This trend suggests that the stress relaxation behavior of DCC-crosslinked hydrogels cannot be accurately predicted by simply applying traditional polymer physics models for entangled melts.

**FIGURE 2:**
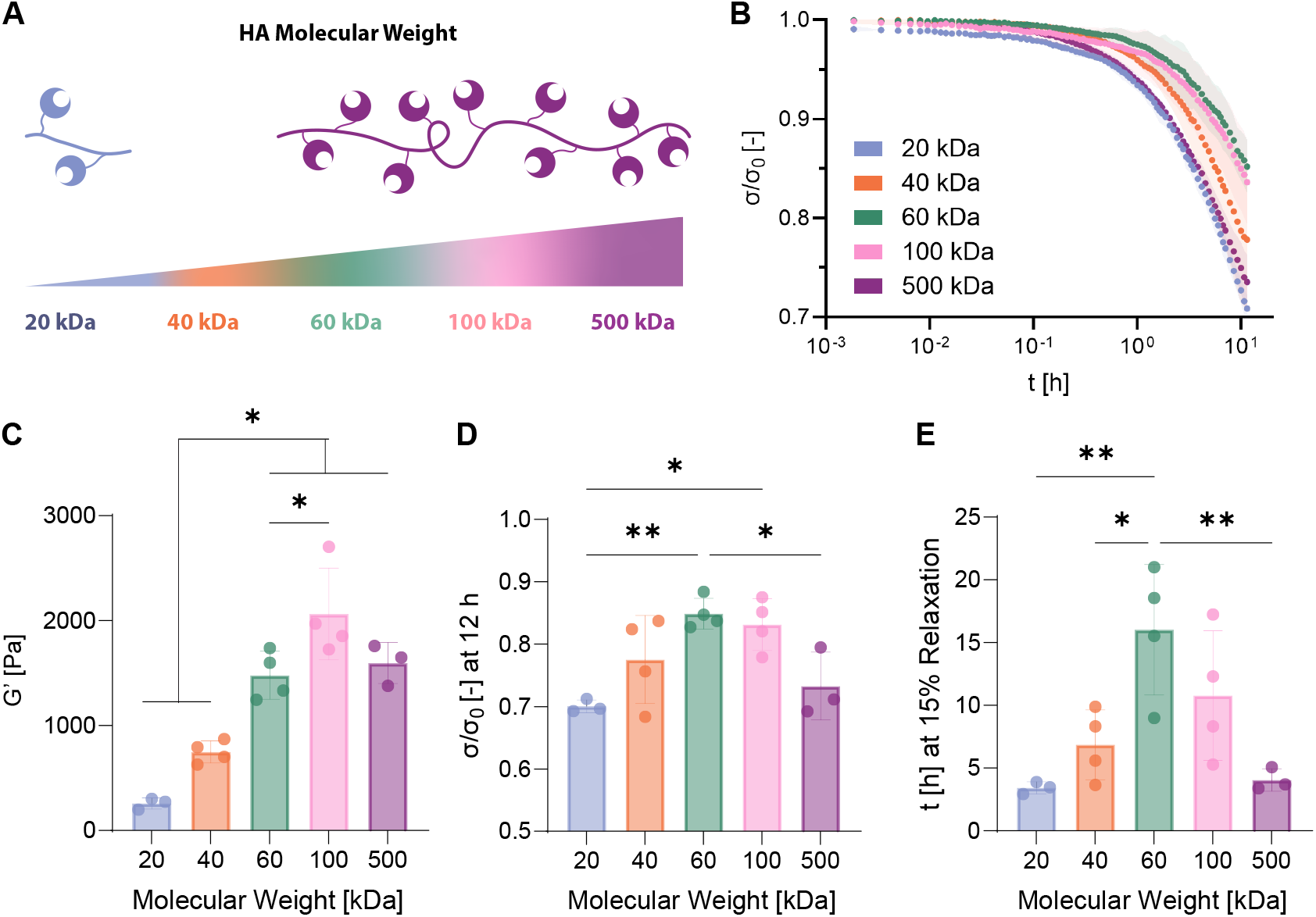
The stiffness and stress-relaxation kinetics of HELP hydrogels is tuned through HA *M*_*w*_. **A**. Schematic and color code of the different HA *M*_*w*_ formulations used in the study. All formulations used HA with 12 % BZA functionalization, Fig. S1. **B**. Fraction of non-dissipated stress (*σ*/*σ*_0_) against time. **C**. Storage modulus (*G′*) at an angular frequency of 1 rad s^−1^. **D**. Fraction of non-dissipated stress after 12 hours. **E**. Time necessary to dissipate 15 % of applied stress. **C-E**. One-way ANOVA, p<0.05 *, p<0.01 **, p<0.001 ***, p<0.0001 ****. Specific p-values are listed on Table S2 (C), Table S3 (D) and Table S4 (E). **B-E**.Averages of *n* ≥ 3 with standard deviation.

### 2.3 Modified Maxwell Modeling of DCC Hydrogels

Given this divergence from the expected stress-relaxation behavior based on polymer melt modeling, we hypothesized that the mismatched sizes of the polymer chains, *i.e*. longer HA (> 100 kDa) and relatively short ELP (*≈* 40 kDa), could be impacting the observed mechanics. Previously, higher polymer *M*_*w*_ has been shown to cause network defects and possible entanglements in the HELP system, which can affect other properties such as erosion profile, stiffness, and injectability de Paiva Narciso et al. (2023). For example, while polymer *M*_*w*_ does not typically impact hydrogel stiffness for irreversibly crosslinked networks, the brachyation theory of reversibly crosslinked polymer networks predicts that as polymer *M*_*w*_ increases, the hydrogel stiffness also increases due to a greater number of possible percolating paths Cai et al. (2022).

By extension, here we propose that the mismatch in chain sizes are leading to different polymer reptationde Gennes (1971) and diffusion regimes that affect the final relaxation profile to a greater extent than the kinetics of the DCC bonds. To explore this idea, we introduce the parameter *q*, which is the ratio of HA to ELP molecular weight. When the HA chain is smaller than the ELP polymer (*i.e*. HA *≈* 20 kDa, ELP *≈* 40 kDa, *q* = 0.5), the chains are unable to span across multiple neighboring ELP chains,de Paiva Narciso et al. (2023), Cai et al. (2020) leading to shorter or no percolating paths, increasing chain mobility, and allowing for faster relaxation (Figure 3A). At matched sizes of HA and ELP polymers (*i.e*. HA *≈* 40 kDa, *q* = 1.0), we envision an ideal network with continuous percolating paths and optimal crosslinking at the stoichiometric ratio of the hydrazone bond (Figure 3B).de Paiva Narciso et al. (2023) When the HA polymer is slightly larger than the ELP polymer (*i.e*. HA *≈* 60-100 kDa, *q* = 1.5 – 2.5), the HA would not only span across neighboring ELP chains and create long percolating paths, but also loop and entangle around the chainsde Paiva Narciso et al. (2023), thus lowering chain mobility (Figure 3C).

**FIGURE 3:**
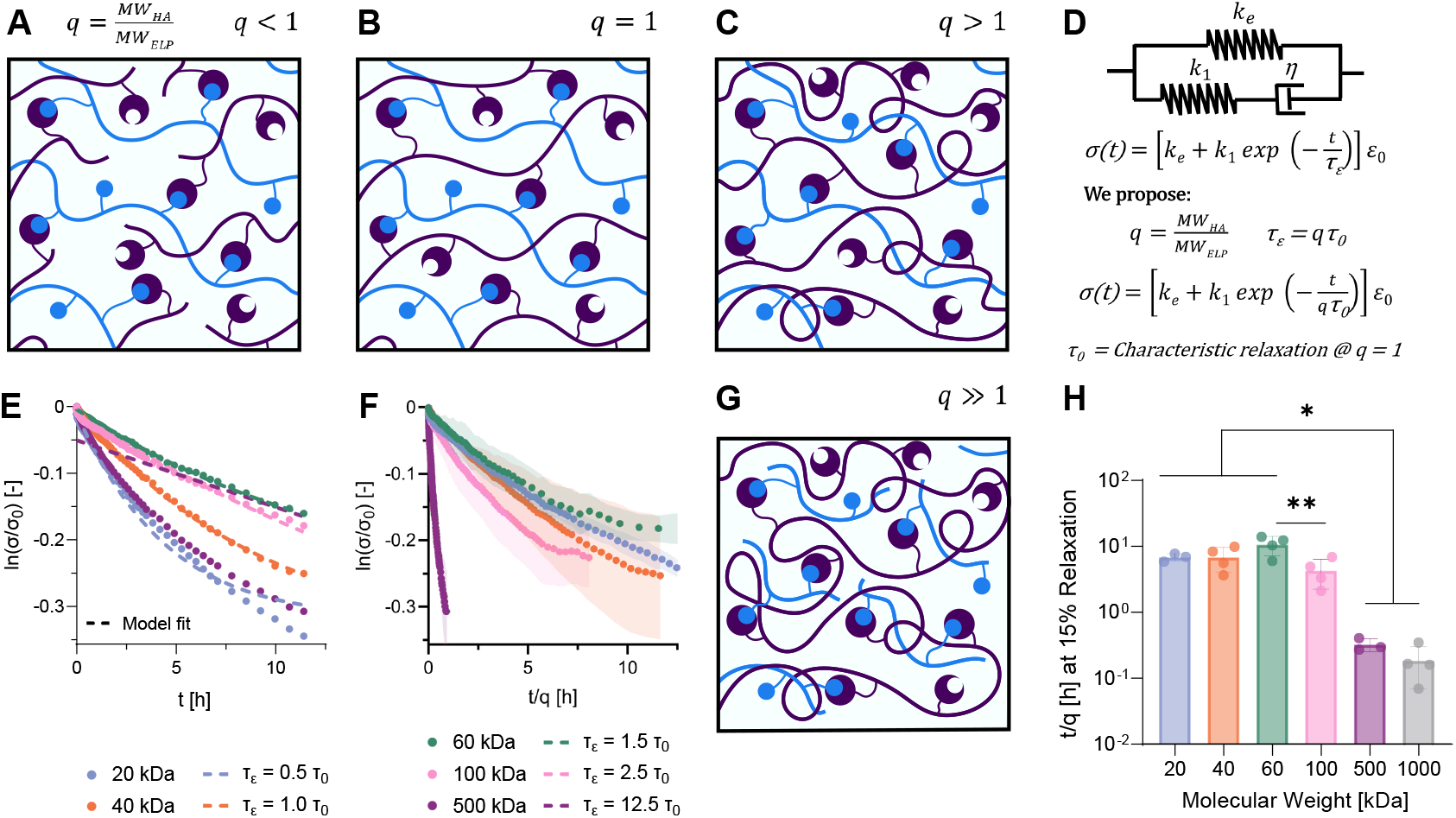
Theoretical framework for the viscoelastic behavior of DCC gels with mismatched *Mw*. **(A-C)**. Schematic of polymer networks formed when mixing ELP-HYD (blue) with HA-BZA (purple) of different *M*_*w*_, where *q* = *MW*_*HA*_/*MW*_*ELP*_. (**A**) When *q* < 1, short percolation paths are formed. (**B**) When *q* = 1, a more uniform continuous network is formed. (**C**) When *q* > 1, a continuous and entangled network forms. **D**. Modified Maxwell Model of viscoelasticity including *q* parameter to account for polymer mismatched sizes. **E**. Natural log of fraction of non-dissipated stress (*σ*/*σ*_0_) against time for all formulations (average of n=3-4), compared to the theoretical fit of Modified Maxwell Model. Fitted values for Modified Maxwell Model are listed on Table S5. **F**. Natural log of fraction of non-dissipated stress (*σ*/*σ*_0_) against time normalized with respect to the q parameter for all formulations (average of n=3-4). **G**. Schematic of polymer networks formed where *q* >> 1, leading to the trapping of the shorter ELP chains within defects and entanglements, preventing them from making effective percolating paths. **H**. Time necessary to dissipate 15 % of applied stress for the different HA *M*_*w*_ HELP formulations normalized with respect to *q* parameter. One-way ANOVA, p<0.05 *, p<0.01 **, p<0.001 ***, p<0.0001 ****. Specific p-values are listed on Table S6.

To investigate this hypothesis, we propose applying a modified version of the Maxwell standard linear solid model commonly used for viscoelastic materials.Chaudhuri (2017), Findley and Davis (2013), Calhoun et al. (2019) In this Modified Maxwell Model, the elastic behavior is represented by a spring, with constant *k*_*e*_, while the viscous part of the material is modeled through a dashpot with viscosity *η*, coupled to a spring with constant *k*_1_, such that the characteristic relaxation time is defined as *τ*_0_ = *η*/*k*_1_ (Figure 3D). In this model, stress relaxation behavior after an applied constant strain is mathematically represented by:

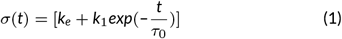

Here, we propose the inclusion of the parameter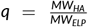count for the mismatched sizes of the polymer chains, such that the characteristic relaxation time is *τ*_*ϵ*_ = *q · τ*_0_, and at *q* = 1, *τ*_*ϵ*_ = *τ*_0_.

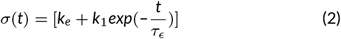

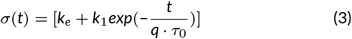

When applying this model to the stress relaxation profiles of our HELP hydrogels, we see that the model achieves good correlation with the experimental data for formulations with HA *M*_*w*_ spanning 20-100 kDa, with *R*2 > 0.98 (Figure 3E, fitting values in Supporting Information, Table S5). This modified model describes the behavior of the relaxation for all formulations with an HA *M*_*w*_ lower than 500-kDa, supporting our hypothesis that the different relaxation profiles are being driven by the mismatched sizes of the polymer chains, whose fitting values can be found in Table S5. Further, when scaling the relaxation behavior with respect to *q*, effectively normalizing and removing the impact of the mismatched polymer sizes, all the different formulations with *MW*_*HA*_ ≤ 100*kDa* collapse into the same relaxation profile (Figure 2F), despite having significantly different stiffness (Figure 2C). This suggests that formulations that fit this model have similar physical processes underlying their relaxation behavior, which is likely the kinetics of the DCC bonds. In addition, as each formulation has the same theoretical number of DCC crosslinks (given the constant functionalization of 12 % BZA for each HA *M*_*w*_), this also implies that the changes in stiffness are likely driven by the mismatched polymer sizes.

However, this modified Maxwell model does not accurately predict the behavior of the higher molecular weight formulation (*e.g*. HA *≈* 500 kDa, *q* = 12.5). We hypothesize that the highly mismatched sizes of the two polymers lead not only to a mildly decreased chain mobility, but to the actual trapping of the shorter ELP chains within defects and entanglements, preventing them from making effective percolating paths (Figure 3G). de Gennes (1971), Rubinstein and Semenov (2001), Rubinstein and Panyukov (2002), **?** In this scenario, we reasoned that a new relaxation mode could dominate, driven by the entanglements of the larger chain rather than the kinetics of the DCC bonds (Figure 3G). To test this hypothesis, we designed a formulation with an even larger HA *M*_*w*_ (*e.g*. HA *≈* 1 MDa, *q* = 25), still functionalized with BZA at 12 %. Interestingly, the 1-MDa formulation also relaxed very rapidly, similar to the rate of the 500-kDa formulation, but displayed a more linear temporal profile, suggesting that the relaxation behavior is primarily governed by entanglements from the onset of relaxation. This is further supported by analyzing the normalized relaxation time (*t*/*q*) to reach 10 % relaxation, which is the same for all formulations with HA *M*_*w*_ ≤ 100 kDa, and drops to a significantly lower value for formulations with HA *M*_*w*_ ≥ 500 kDa (Figure 3H). Thus, as an alternative strategy to selecting different DCC crosslinking reactions to control the hydrogel stress-relaxation rate, we demonstrate that tuning the molecular weight of one of the components (*i.e*. thereby altering *q*) can significantly impact network relaxation dynamics.

### 2.4 Impact of HA degree of functionalization on stiffness and stress-relaxation kinetics

Having explored the impact of HA:ELP *M*_*w*_ on stress relaxation, we set out to investigate the effect of the degree of chemical functionalization of HA. We first tested functionalization at 6%, 12%, and 20% BZA for the 20 kDa and 60 kDa HA *M*_*w*_ formulations, corresponding to stoichiometric ratios (*r*) of BZA:HYD reactive groups of 0.41, 0.81, and 1.35, respectively (Figure 4A-D). Here, it is important to highlight that at *r* = 1, the concentration of available reactive groups will match the stoichiometry of the hydrazone bond reaction, which would be expected to produce the stiffest hydrogel in a traditional, static covalent crosslinked system (Figure 4C). However, for DCC chemistry, the crosslinking reaction has an equilibrium constant (*k*_*eq*_), which controls the number of on- and off-bonds. McKinnon et al. (2014) Thus, by increasing the concentration of BZA (*r* > 1), we favor the formation of new bonds, increasing the amount of crosslinks formed at any given time within the network (Figure 4D). In general, this is supported by our data, where the increased functionalization leads to increased stiffness within the same family of HA *M*_*w*_ hydrogels, attributed to a higher crosslinking density (Figure 4E). In contrast, when *r* < 1, the low amount of crosslinks leads to lower stiffness. This is particularly evident in the 20-kDa HA functionalized with 6% BZA, where no hydrogel is formed, likely due to the combined effect of low functionalization and short HA chains leading to the absence of a percolating network (Figure 4B,E). Thus, the degree of functionalization is an experimental variable that can be used to alter the network connectivity, and hence stiffness, in DCC-crosslinked hydrogels.

**FIGURE 4:**
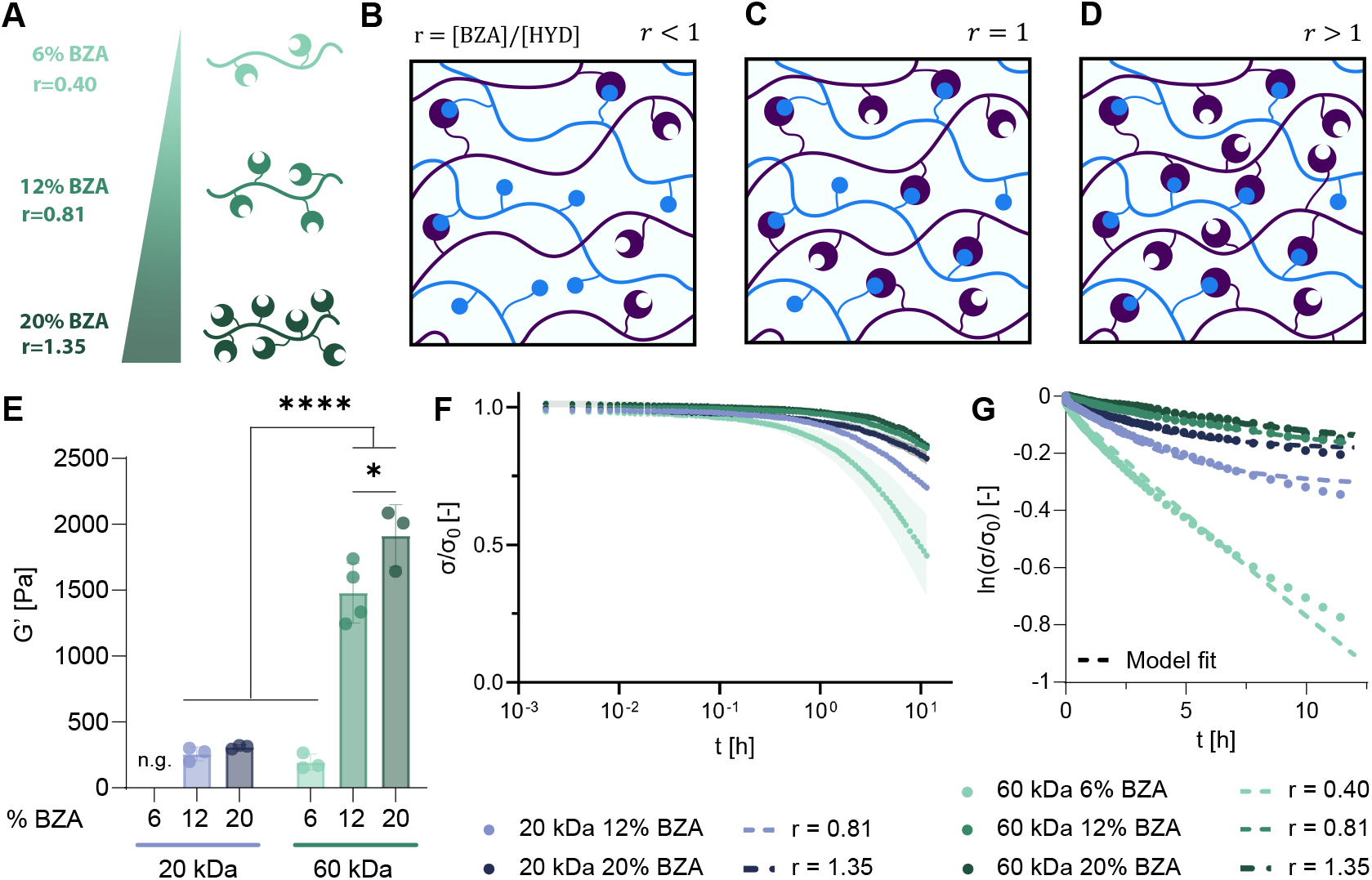
The stiffness and stress-relaxation of HELP gels is tuned through the degree of functionalization of HA. **A**. Schematic and color code of the different HA degrees of functionalization (Fig.S1) used in the study, resulting in different ratios of the DCC reactive groups (*r* = [*BZA*]/[*HYD*]). **B-D**. Idealized schematics of polymer networks formed when mixing ELP-HYD (blue) with HA-BZA (purple) with different degrees of functionalization, resulting in *r* < 1 (B), *r* = 1 (C), and *r* > 1 (D). **E**. Storage modulus, *G′*, for the different HELP formulations. One-way ANOVA, p<0.001 ***, p<0.0001 ****. Specific p-values are listed on Table S7. **F**. Fraction of non-dissipated stress (*σ*/*σ*_0_) against time. **G**.Natural log of fraction of non-dissipated stress (*σ*/*σ*_0_) against time for formulations compared to the theoretical fit of Modified Maxwell Model. Fitted values for Modified Maxwell Model are listed on Table S5. **E-G**. Averages of *n* ≥ 3 with standard deviation.

We reasoned that these alterations in network connectivity would also offer a means to tune the stress-relaxation rate. Consistent with this idea, at both lower and intermediate HA *M*_*w*_ (*i.e*. 20 and 60 kDa), the hydrogels exhibited changes in stress-relaxation rate. Higher crosslinking density led to decreased chain mobility, with fewer degrees of freedom and slower relaxation, as previously reported in literature (Figure 4F,G). This behavior is observed at low HA *M*_*w*_, with the 12 % BZA formulation relaxing faster than the 20 % BZA, as well as at intermediate HA *M*_*w*_, with the 12 % BZA formulation relaxing significantly slower than its 6 % counterpart. Interestingly, there seems to be a plateau to the contribution of crosslinking density in slowing down relaxation, as the 60-kDa HELP gels with 12 % and 20 % BZA functionalization have overlapping relaxation profiles. Importantly, our modified Maxwell model accurately describes the stress-relaxation behavior for these five HELP formulations (Figure 4G), with the decrease in crosslinking density captured by the decrease in the *k*_*e*_ (Table S5, Fig. S3) value for hydrogels with lower degree of functionalization. Thus, while keeping the identity of the DCC chemistry the same (*i.e*. all BZA-HYD crosslinks), the stress-relaxation rate of the hydrogel can be tuned either by changing polymer molecular weight or by changing the degree of functionalization.

### 2.5 Modified Maxwell model for large polymer chains

Interestingly, for HELP gels with relatively high HA *M*_*w*_ (≥ 500 kDa) resulting in large HA:ELP *M*_*w*_ mismatch with *q* >> 1, the degree of HA functionalization impacts gel mechanics in a different way compared to formulations with *q* near 1.

In particular, the stiffness decreases considerably as HA functionalization is increased from 12% to 20% (*i.e. r* from 0.81 to 1.35) (Figure 5A). This supports our hypothesis that the large *M*_*w*_ mismatch is leading to the formation of a network with more entanglements and defects that prevent effective crosslinking (Figure 5B-D).de Gennes (1971), Ma et al. (2025), **?** Consistent with this idea, as q is increased even further (HA *M*_*w*_ 1 MDa, *q* = 25) while keeping the degree of functionalization constant (12 %, *r* = 0.81), the resulting gel stiffness continues to decrease (Figure 5A). To further highlight the increased potential/probability to form entanglements as HA *M*_*w*_ increases, we calculated the theoretical entanglement concentration for unmodified HA in solution with a good solvent (Table S9, Supplemental Methods).Cai et al. (2022), de Paiva Narciso et al. (2023), Oelschlaeger et al. (2013), Rubinstein et al. (2003) For HA *M*_*w*_ of 500 kDa and 1 MDa, the theoretical entanglement concentrations are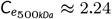*wt*% and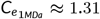*wt*%, respectively, which are similar to or lower than the concentration of our precursor HA-BZA solution (2 *wt*%). Thus, as the HA *M*_*w*_ increases, homogenous mixing of the two polymeric solutions becomes more challenging, resulting in uneven network formation with defects. This prevents the formation of a uniformly crosslinked network and leads to low effective crosslinking density and lower stiffness (Figure 5A). As with the intermediate HA *M*_*w*_ formulations, the higher amount of entanglements and decreased chain mobility for higher HA functionalization are also reflected in the observed stress-relaxation behavior. Similar to formulations with intermediate *M*_*w*_, lower functionalization leads to a higher stress relaxation rate that decreases with increasing functionalization up to a plateau value (Figure 5E).

**FIGURE 5:**
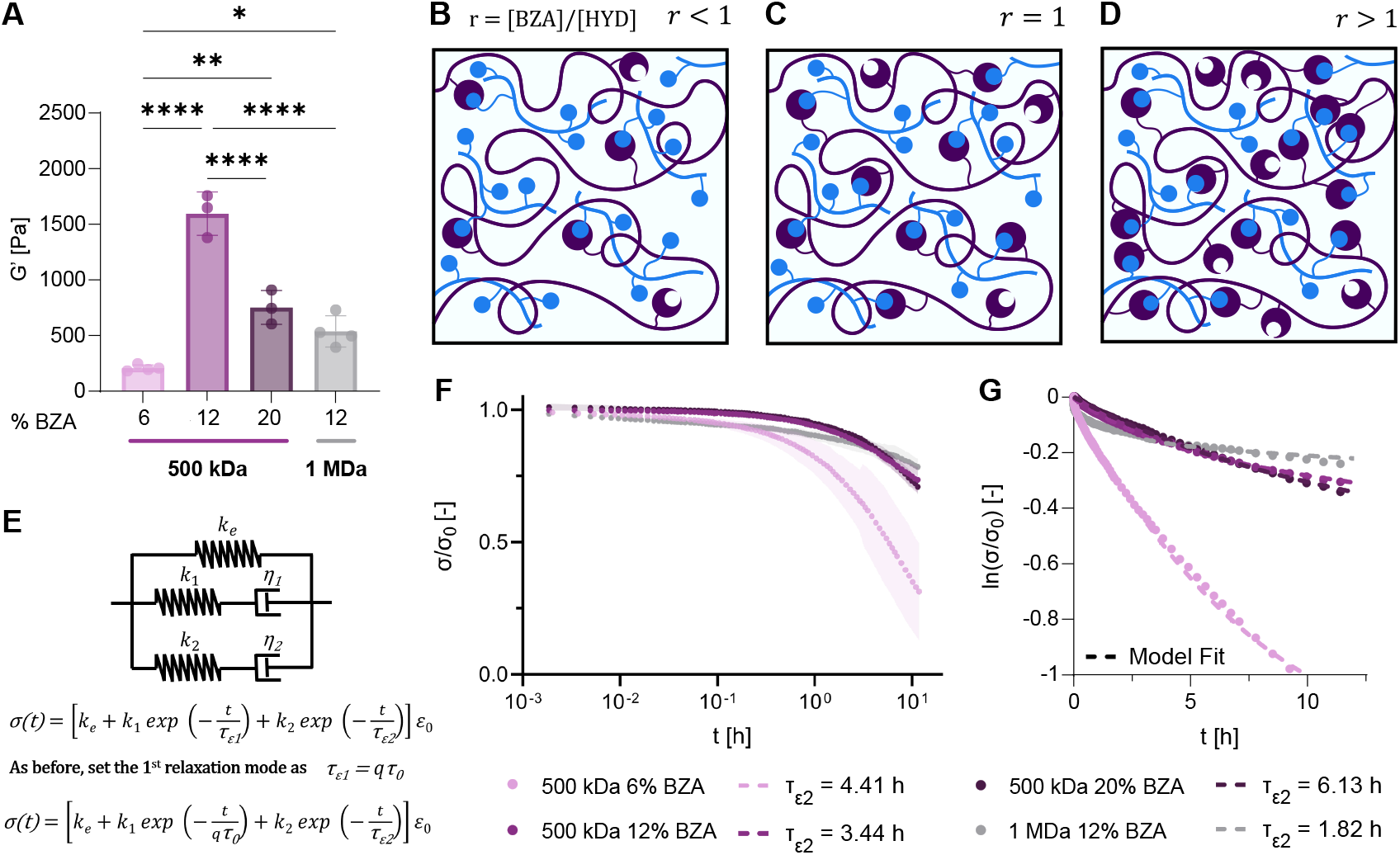
The stress-relaxation in large mismatched sizes is driven both by entanglements and DCC kinetics. **A**. Storage modulus *G′* for the different HELP formulations with high HA *M*_*w*_ . One-way ANOVA, p<0.001 ***, p<0.0001 ****. Specific p-values are listed on Table S8. **B-D**. Schematic of polymer networks formed when mixing ELP-HYD (blue) with HA-BZA (purple) with different % functionalization, where *r* = [*BZA*]/[*HYD*]. resulting in *r* < 1 (B), *r* = 1 (C), and *r* > 1 (D) **E**. Two-Component DCC Network model of viscoelasticity, including both the *q* parameter to account for polymer mismatched sizes, and a second dashpot representing the additional relaxation mode. **F**. Fraction of non-dissipated stress (*σ*/*σ*_0_) against time. **G**. Natural log of fraction of non-dissipated stress (*σ*/*σ*_0_) against time for formulations (average of *n* = 3 – 4), compared to the theoretical fit of Two-Component DCC Network model. Fitted values for Two-Component DCC Network model are listed on Table S10. Averages of *n* ≥ 3 with standard deviation.

However, as described earlier, our single-dashpot Maxwell model modified with *q* terms still does not accurately fit experimental data for very large *q* (Figure 3E). We reasoned that this was likely due to the larger number of entanglements present in the longer HA chains, which would also contribute a relaxation timescale to the overall network viscoelasticity. Here, to capture this extra relaxation mode, we propose an additional modification to our modified Maxwell model, adding a second spring-dashpot to represent entanglement relaxation, with spring constant *k*_2_ and viscosity *η*_2_, such that the characteristic relaxation time is defined as *τ*_*ϵ*2_ = *η*_2_/*k*_2_, Figure 5E. In this Two-Polymer DCC Network model, the stress relaxation behavior after an applied constant strain is mathematically represented by:

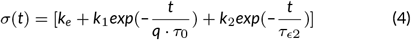

With this further modification, the Two-Polymer DCC Network model accurately captures the viscoelastic behavior of all formulations tested, including the high q formulations (*i.e*. HA *M*_*w*_ 500 kDa and 1 MDa) (Figure 5G, Fig. S2, Table S10 and Fig. S3).

This suggests that at low functionalization, the relaxation is being dominated by the movement of the long uncrosslinked entangled chains, leading to a faster rate of relaxation (Figure 5B). As we increase the functionalization, the kinetics of the bonds slow down the relaxation (Figure 5C), as the crosslinking density increases. However, at higher functionalization, there is a notably lower crosslinking density as reflected in the decrease in stiffness. Thus, this suggests that the slower stress relaxation seen at high functionalization in high HA *M*_*w*_ formulations is likely also dominated by the movement of entangled chains, which have highly restricted mobility due to entrapment of the ELP chains (Figure 5D). This decrease in mobility is also captured by our model, as while *τ*_*ϵ*2_ has similar values for our 6 and 12 % BZA formulations of 500 kDa HA, it almost doubles for our 20 %BZA formulation.

### 2.6 Hydrogel viscoelasticity alters neurite morphology

After establishing these design rules for how molecular-level polymer parameters control the stiffness and stress-relaxation kinetics of HELP hydrogels, we next evaluated the effect of gel mechanics on cellular behavior. Specifically, we explored how gel viscoleasticity impacted the morphology and neurite outgrowth from human-induced pluripotent stem cell-derived neural progenitor cells (hiPSC-NPCs), as these cells are known to be exquisitely mechanosensitive.Huang et al. (2025b), **?**), Navarro et al. (2022) There are inherent difficulties associated with directly studying the human brain, as samples are restricted to postmortem and biopsy tissues, providing limited information since they are constrained to a specific timepoint in neural development and disease progression.Stiles and Jernigan (2010), Liu et al. (2023), Calhoun et al. (2019) In a similar manner, the information acquired by functional neuroimaging is also limited due to inherent resolution challenges. Kalin (2021) Moreover, the applicability of *in vivo* models developed from studying rodent brains is restricted by the anatomical and physiological disparities between the two.Chesselet and Carmichael (2012), Eliot and Richardson (2016) Therefore, the advancement of human *in vitro* models are critically important to study human brain development and disease. To probe the mechanical versatility of our HELP gel formulations and their application in 3D *in vitro* tissue culture models, we encapsulated hiPSC-NPCs in HELP gels of five different formulations (Figure 6).

**FIGURE 6:**
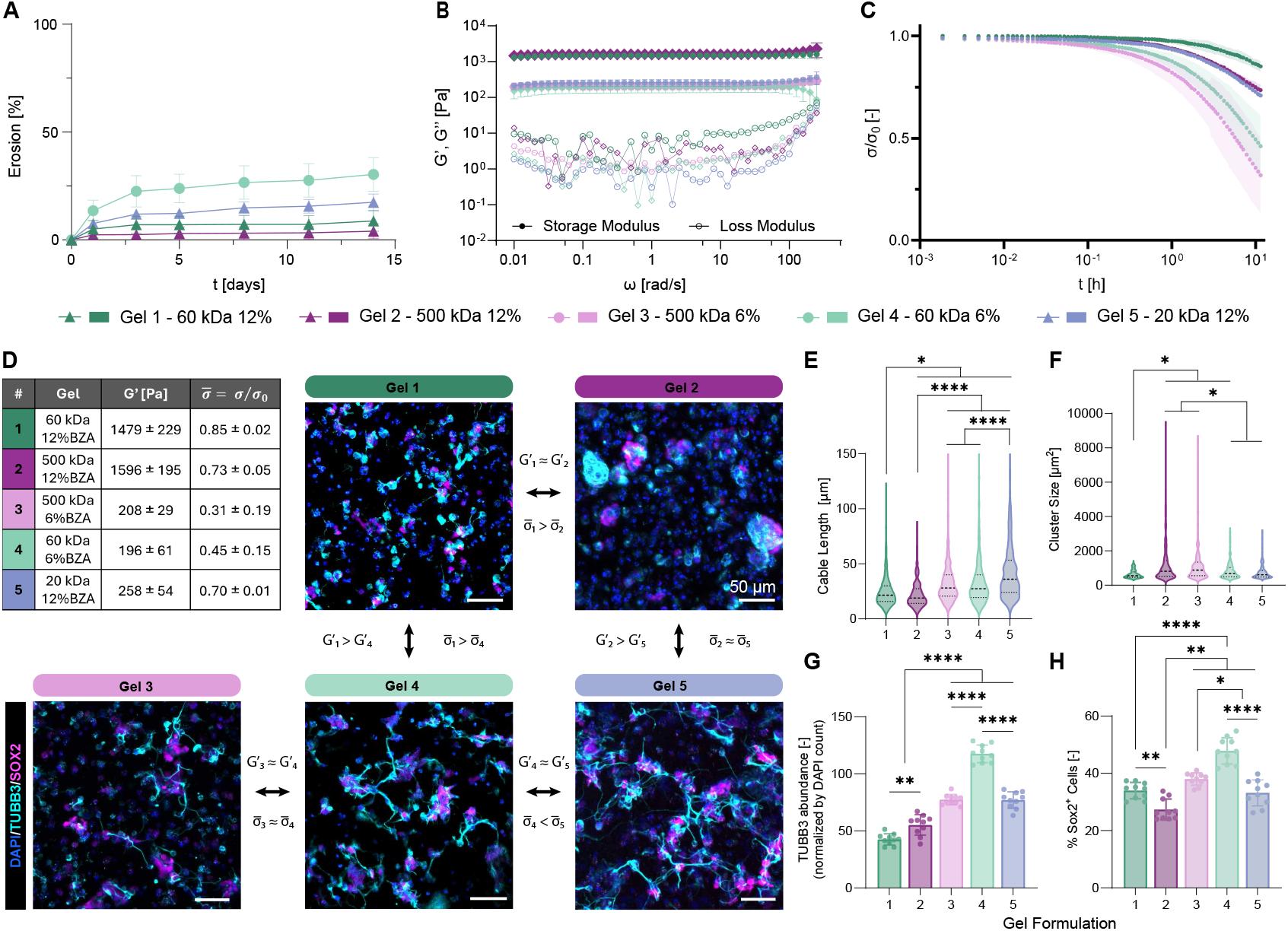
The viscoelastic characteristics of hydrogels impacts neurite morphology in 3D tissue culture system. **A**. Erosion profile of selected HELP gels. **B**. Frequency sweep of selected HELP gel formulations ranging from 0.01 to 250 rad/s, storage modulus (G’, filled symbols) and loss modulus (G”, open symbols).**C**. Fraction of non-dissipated stress (*σ*/*σ*_0_) against time. **D**. Table with selected HELP gel formulations with their respective storage modulus (G’), and fraction of non-dissipated stress at 12 hours (*σ* = *σ*/*σ*_0_), Top right. Representative maximum projection fluorescence images of hNPCs encapsulated at 30,000 cells/μl within all five gel formulations after 7 days of culture with labeled cell nuclei (DAPI, blue) and neuritic extensions (TUBB3, cyan), and stemness marker (Sox2, magenta), with comparison relationships. **E**. Cable length of individual neurites through quantifying TUBB3-expressing projections with the SNT toolbox. **F**. Cell-cluster sizes wuantified based on DAPI staining, identifying objects > 40 pixel units in diameter. Kruskal-Wallis, p<0.05 *, p<0.0001 ****. Specific p-values are listed on Table S11(E) and Table S12(F). **G**. Tubulin abundance normalized by nuclei (DAPI) count. **H**. Percentage of Sox2+ cells. One-way ANOVA, p<0.05 *, p<0.01 **, p<0.0001 ****. Specific p-values are listed on Table S13(G) and Table S14(H).

One of the key advantages of controlling viscoelasticity through our chosen molecular parameters, HA *M*_*w*_ and degree of functionalization with BZA, is the increase in long-term hydrogel stability (Figure 6A). Standard approaches to tune stress-relaxation by altering the DCC bond kinetics, such as incorporating ALD moieties as the crosslinking motif, have been shown to result in rapid hydrogel erosion (Figure 1G). Our novel approach retains the use of BZA moieties, known to have a slower on-off binding rate. This leads to a decrease in erosion for all selected gel formulations, with a maximum of 25 % erosion over two weeks (Figure 6A), while ALD formulations experienced 75-100% over the same time period (Figure 1G). We selected the HELP gel formulations such that we would have a set of stiffness-matched gels and a set of stress-relaxation-matched gels (Figure 6B-C), allowing us to separately investigate the effects of each variable. The hydrogels selected for the *in vitro* study had the following HA *M*_*w*_ and degree of functionalization parameters: 60 kDa and 12 % BZA (Gel 1), 500 kDa and 12 % BZA (Gel 2), 500 kDa and 6 % BZA (Gel 3), 60 kDa and 6 % BZA (Gel 4), and 20 kDa and 12 % BZA (Gel 5) (Figure 6D). Briefly, Gel 1 and Gel 2 have similar stiffness, while Gel 1 has a slower stress-relaxation rate. Gel 2 and Gel 5 have a similar stress-relaxation rate, but Gel 2 has a higher stiffness. Gel 1 has a higher stiffness and a slower stress-relaxation rate than Gel 4. Gel 4 and Gel 5 have similar stiffness, but Gel 5 has a slower stress-relaxation rate. Finally, Gel 4 and Gel 3 have similar stiffness and stress-relaxation rate.

Encapsulated cells were exposed to identical media for 7 days and then immunostained for βIII-tubulin (TUBB3), a neuronal lineage marker and key component of neuritic projections, and Sox2, a marker of NPC stemness (Figure 6D).Roth et al. (2023), Huang et al. (2025b) Quantitative morphology metrics included measurements of cable length to compare neurite extension (Figure 6E) and cell cluster size to compare the propensity to form neurospheres (Figure 6F). NPCs frequently grow as neurospheres when cultured in suspension, and when encapsulated can grow either as neurospheres or dispersed cells.Huang et al. (2025b) In our earlier work, we have shown that hiPSC-NPCs encapsulated in fast stress-relaxing, intermediate stiffness (*G′ ≈* 800 Pa) HELP gels using a 50:50 BZA:ALD ratio (12 % functionalization) exhibited a highly branched morphology with neurites extending throughout the gel from dispersed cells.Roth et al. (2023)

When comparing Gel 1 and Gel 2 (*G′*1 *≈ G′*2 and *σ*_1_ > *σ*_2_), while the faster relaxation rate did not lead to increased neurite outgrowth, there was significantly higher expression of *β*III-tubulin (TUBB3), Figure 6E-H. The expression of neural stemness marker Sox2 was higher in Gel 1, with a higher number of Sox2+ cells. There was also the presence of large cell clusters in Gel 2 (HA 500 kDa 12 %BZA formulation), which appeared to be where the majority of *β*III-tubulin expression was localized. Looking at Gel 2 and Gel 5 (*G′*2 > *G′*5 and *σ*_2_ *≈ σ*_5_), these similar relaxation rates led to widely differing results, suggesting this phenomena may be driven by the lower stiffness of Gel 5, which resulted in increased neurite outgrowth, smaller cell clusters, and higher expression of both βIII-tubulin and Sox2. We saw similar behavior when comparing the cellular responses in Gel 1 and Gel 4 (*G′*1 > *G′*4 and *σ*_1_ > *σ*_4_), with the lower stiffness corresponding to longer neurites with higher expression of βIII-tubulin and Sox2. For Gel 4 and Gel 5 (*G′*4 *≈ G′*5 and *σ*_4_ < *σ*_5_), which are both of lower stiffness, the faster relaxation rate of Gel 4 corresponded to a higher expression of βIII-tubulin and Sox2, although the cable length, our metric for neurite outgrowth, was higher for Gel 5 (HA 20 kDa 12 %BZA formulation), suggesting that there may be an optimal combination of stiffness and relaxation rate to promote neurite extension. An unexpected finding is that Gel 2, with intermediate relaxation rate (*σ*_1_ > *σ*_2_ > *σ*_4_) and high stiffness, presented the lowest cable length of all conditions. This formulation (Gel 2) also led to the formation of large cell clusters with high expression of βIII-tubulin (Figure 6F).

These data may indicate that the higher HA *M*_*w*_ and the secondary relaxation mode driven by entanglements might hinder neurite outgrowth. One way to investigate this is to compare the formulations with similar stiffness and stress-relaxation rate, Gel 4 and Gel 3 (*G′*4 *≈ G′*3 and *σ*_4_ *≈ σ*_3_), but differing HA *M*_*w*_ (60 kDa and 500 kDa, respectively) (Figure 6F). Although they had similar cable lengths, the lower HA *M*_*w*_ formulation (Gel 4) resulted in higher expression of both βIII-tubulin and Sox2 and fewer cell clusters (Figure 6G,H). These data are consistent with the idea that higher HA *M*_*w*_ may impact cell morphology, potentially through secondary relaxation modes caused by polymer entanglements, leading to the formation of cell clusters. We hypothesize this behavior could be due to the lower crosslink concentration, which may lead to local relaxation rather than continuous bulk matrix relaxation.

Overall, these results provide insight into the potential effects of both stiffness and stress-relaxation on cell morphology, agreeing with previous results and demonstrating that the synergy of lower stiffness with fast stress-relaxation rates can correlate with increased stemness maintenance and neurite outgrowth.Huang et al. (2025b), Navarro et al. (2022), Roth et al. (2023) Further, the enhanced stability of these formulations lend themselves to future cellular biomechanics studies to explore in detail the mechanosignaling pathways responsible for the observed phenotypes.

## 3 CONCLUSIONS

Three-dimensional models of *in vitro* tissue culture enable high-throughput drug screening, mechanistic studies, and the investigation of human development, making them indispensable for biomedical research. Further, their ability to recapitulate biochemical and mechanical characteristics of tissues allows the field to move away from animal studies and to explore *in vitro* models as animal-free “new approach methodologies” (*i.e*. NAMs). The use of hydrogels as 3D tissue culture models has been widely explored, as their tunable viscoelastic and biochemical properties are known to impact cell morphology and function. Particularly, DCC croslinked gels have risen to prominence since the reversible nature of their bonds allows for the introduction of stress-relaxation, a key mechanical characteristic of native ECM. Traditional methods to tune the stress-relaxation and viscoelasticity of DCC gels are based on bond kinetics, which directly affect gel erosion rates, typically preventing their use for multi-day 3D tissue culture. Here, we demonstrate the use of molecular parameters (HA *M*_*w*_ and degree of BZA functionalization) to control the viscoelasticity of DCC hydrogels while still allowing for week-long culture experiments.

Specifically, by varying only the HA *M*_*w*_ in our HELP system, we investigated the effects of mismatched polymer sizes (HA vs. ELP) on gel stress relaxation and stiffness behavior. As expected, higher molecular weight typically led to higher gel stiffness (*G′*). Longer polymers have a greater number of potential crosslinking sites per chain, and hence a larger number of possible percolating paths through the network Cai et al. (2022). Interestingly, we observed that the mismatched polymer sizes strongly influence the network stress-relaxation rates. Varying HA *M*_*w*_ had a parabolic effect on the relaxation rate of the gels: formulations with high HA *M*_*w*_ (500 kDa) relaxed as fast as low HA *M*_*W*_ (20 kDa). To understand this, we developed and validated a modified Mawell Model of gel viscoelasticity, in which we include the ratio of polymer molecular weight, *q* = *MW*_*HA*_/*MW*_*ELP*_, as a normalizing parameter. This model fits the behavior of all formulations with low and intermediate HA *M*_*w*_, where their relaxation behavior collapses into a single trendline when normalized by *q*. However, this model fails to replicate the behavior of the high HA *M*_*w*_ formulation (500 kDa), suggesting additional mechanisms of relaxation are in place when the polymers are highly mismatched in size, likely driven by entanglement movement. The relaxation of HELP gels is further tunable through the degree of functionalization of BZA on our HA chain, with increasing functionalization leading to slower-relaxing hydrogels due to the increase in crosslinking density. While our modified Maxwell model captures this increase in elasticity in low and intermediate HA *M*_*w*_, it again fails to explain the behavior at very high HA *M*_*w*_ (≥ 500 *kDa*). To address this, we introduced the Two-Component DCC Network model, where, in addition to the *q* parameter, we added a second spring-dashpot to represent a secondary mode of relaxation. This theoretical framework was able to describe the viscoelastic behavior of all characterized HELP gel formulations across the full ranges of HA *M*_*w*_ and degree of functionalization that were tested. Thus, this model provides a helpful conceptual understanding in which to predict the viscoelastic behavior of HELP gels. As DCC gels can be designed using a variety of different polymer backbones and crosslink chemistries, an exciting area for future study is to explore if the Two-Component DCC Network model developed here can also describe the viscoelastic behavior of other DCC gels. For example, DCC hydrogels with alternative kinetics, such as oxime and imine bonds, as well as alternative polymer backbones, such as alginate or polyethylene glycol, can be studied to explore the applicability of the theoretical framework described here.

Given the importance of gels as 3D *in vitro* tissue culture systems, we then demonstrated the applicability of these formulations to culture hiPSC-NPC for one week. As a key advantage, we were able to keep the crosslinking chemistry (and hence crosslinking reaction kinetics) constant across all formulations, resulting in gels that were sufficiently stable for multi-day culture. By separately modulating the stiffness and relaxation rate, we observed that softer gels (*G′ ≈*200 Pa) with intermediate to fast stress-relaxation rates lead to higher expression of a stemness marker and increased neurite outgrowth. Interestingly, we also observed that gels formulated with high HA *M*_*w*_ (500 kDa) correlated with increased formation of cell clusters (*i.e*. neurospheres), suggesting that secondary relaxation modes may also influence cell phenotype. Cells can “sense” the viscoelastic properties of their surrounding matrix through several different mechanisms; thus, this family of modular gels is well suited for future studies of the mechanisms of cellular mechanosignaling. Chaudhuri et al. (2020), Suhar et al. (2023), Huang et al. (2025b)

In summary, this work expands the molecular design space of DCC hydrogels to achieve tunable viscoelasticity for 3D *in vitro* models, decreasing our reliance on bond kinetics and improving our conceptual understanding of relaxation modes. Given the large design space of DCC hydrogels and their ability to support long-term culture of human cells, these materials are well suited to replace and complement animal studies of disease progression and drug discovery in the future.

## 4 EXPERIMENTAL

### 4.1 Materials

All reagents were purchased from Sigma-Aldrich and used without further purification, unless otherwise noticed.

#### 4.1.1 Synthesis of Elastin-Like Protein

The ELP was synthesized through recombinant protein expression as previously described de Paiva Narciso et al. (2023). Briefly, ELP encoding plasmids were transformed into BL21(DE3) pLysS *Escherichia coli* (Life Technologies). The bacteria were grown in Terrific Broth at 37 *°*C, up to an OD600 of 0.8, at which point ELP expression was induced by isopropyl *β*-d-1-thiogalactopyranoside (Fisher), at a final concentration of 1 mM . Expression proceeded for 7 h, at which point the bacteria were pelleted and resuspended in TEN Buffer (10 mM Tris (Fisher), 1 mM ethylenediamine tetraacetic acid disodium salt dihydrate (EDTA, Fisher), and 100 mM NaCl (Fisher), pH 8.0). de Paiva Narciso et al. (2023). The cell-pellet was lysed through alternating freeze and thaw cycles. Deoxyribonuclease and phenylmethylsulphonyl fluoride were added to the cell lysate at 1 mM. de Paiva Narciso et al. (2023). The resulting ELP was purified through thermocycling, dialyzed in deionized water (DI) for three days at 4 *°*C, then frozen, and lyophilized. de Paiva Narciso et al. (2023). Final lyophilized material was stored at −20 *°*C.

Functionalization and Characterization of ELP-HYD The ELP was modified following previously published protocols,LeSavage et al. (2018), Hefferon et al. (2023) by modifying the amine on the lysine amino acid through a hexafluorophosphate azabenzotriazole tetramethyl uronium (HATU) enabled amidation reaction. In summary, the lyophilized ELP was dissolved in a 1:1 solution of anhydrous dimethyl sulfoxide (DMSO) and *N,N*-dimethylformamide (DMF) at a 3% (w/v). de Paiva Narciso et al. (2023)In a separate vessel at room temperature, tri-boc hydrazinoacetic acid was dissolved at 2.1% (w/v) in anhydrous DMF. After dissolution, the tri-boc was activated with HATU (2.0 equivalence per ELP amine; Sigma, 445460) and 4-methylmorpholine (NMM, 5 equivalence per ELP amine) for 10 min. de Paiva Narciso et al. (2023)The activated tri-boc molecule was added to the ELP solution dropwise under continuous stirring, and the reaction proceeded overnight.de Paiva Narciso et al. (2023) The tri-boc modified protein was precipitated in ice-cold diethyl ether in a 1:4 volume ratio of reaction solution to ether. de Paiva Narciso et al. (2023)The product was pelleted and dried under nitrogen for 3 days.de Paiva Narciso et al. (2023) Once dried, the pellets were resuspended in a 1:1 solution of dichloromethane (DCM) and trifluoroacetic acid with 5% (v/v) tri-isopropylsilane (Sigma, 233781) for 4 h to deprotect the hydrazine group. The deprotected ELP-HYD was precipitated in ether, dried under nitrogen for 3 days, and resuspended in DI water at 4 *°*C. de Paiva Narciso et al. (2023)The protein was dialyzed against DI water for three days at 4 *°*C. The resulting solution was sterile filtered (0.22 *µ*m) in a biosafety cabinet, lyophilized, and stored at −20 *°*C.

### 4.1.2 Functionalization and Characterization of Hyaluronic Acid with Benzaldehyde (HA-BZA)

HA was functionalized following previously published protocols.de Paiva Narciso et al. (2023) First, HA with different molecular weight (LifeCore, HA20K-1, HA40K-1, HA60K-1, HA100K-1, HA500K-1, HA1M-1) was modified with an alkyne group through an EDC/NHS reaction at different equivalences depending upon the desired final BZA functionalization (1 eq. for 6% and 12% BZA functionalization, 3 eq. for 20% BZA functionalization).de Paiva Narciso et al. (2023) The reaction proceeded at room temperature overnight. The final mixture was dialyzed for three days against DI water. After dialysis, the mixture was filtered through a 0.22 *µ*m filter and subsequently lyophilized.

To achieve HA-BZA, a copper-catalized azide-alkyne cycloaddition, also known as a copper-click reaction was performed on the HA–alkyne. Briefly, the HA-alkyne solid was dissolved in isotonic 10× PBS (10× iPBS) (81 mM sodium phosphate dibasic, 19 mm sodium phosphate monobasic, 60 mM sodium chloride in Milli-Q water; pH adjusted to 7.4; 0.22 *µ*m filtered) supplemented with *β*-cyclodextrin at a 1 mg mL^−1^. de Paiva Narciso et al. (2023) Stock solutions of 2.4 mm copper sulfate (Copper sulfate, 5-hydrate – JT Baker 1841-01) and 45.2 mm sodium ascorbate were prepared. de Paiva Narciso et al. (2023) All solutions were degassed for 30 min under nitrogen flow. Finally, azido-BZA (Santa Cruz Biotechnology) was dissolved in extra dry DMSO (Acros Organics 61097-1000) depending on the targeted BZA functionalization: 1 eq. with respect to the corresponding HA–alkyne for 6% BZA functionalization, and 2 eq. for 12% and 20% BZA functionalization.de Paiva Narciso et al. (2023) The reagents were added to the solution as previously described, and the reaction proceeded for 24 h at room temperature.de Paiva Narciso et al. (2023) Then, the reaction stopped by chelating the copper with 50 mM EDTA (Fisher O2793-500). de Paiva Narciso et al. (2023) Last, the mixture was dialyzed, filtered, and lyophilized as previously described. The degree of functionalization of HA-BZA was evaluated through 1H NMR,[21, 33] and the different degrees of functionalization were quantified, resulting in (6% *±* 3%, 12% *±* 3%, 20% *±* 3%) (Figures S1, Supporting Information).de Paiva Narciso et al. (2023)

### 4.1.3 Hydrogel Formation

All HELP hydrogels have a final composition of 1 wt% ELP-HYD and 1 wt% HA-BZA in 10× iPBS, regardless of the HA molecular weight or degree of functionalization. The components, HA-BZA and ELP-HYD, were dissolved at a stock concentration of 2 wt% in 10× iPBS overnight at 4 *°*C under continuous rotation. Equal volumes of 2 wt% HA-BZA and 2 wt% ELP-HYD were mixed on ice, to prevent ELP aggregation. This procedure allowed for the hydrazone crosslinks to spontaneously form, creating gels with final concentration of 1 wt% HA and 1 wt% ELP.

### 4.2 Rheometry

Rheological characterization was performed on an ARG2 rheometer (TA Instruments) equipped with a Peltier plate using a 20-mm cone-plate geometry with an angle of 1 *°*. The two components of the HELP hydrogels (ELP-HYD and HA-ALD or HA-BZA) were pre-mixed in an 0.6 mL Eppendorf tube and loaded onto the rheometer with the measurement starting 10 s after mixing. The gelation was monitored through a time-oscillation measurement at an angular frequency (*ω*) of 1 rad s^1–^ and a shear strain amplitude (*γ*) of 1 %. The gel is loaded on the rheometer and kept at 4 *°*C for 5 min. Subsequently the temperature is ramped to 23 *°*C at a heating rate of 3 *°*C min^−1^ and the measurement continues isothermal for 15 min. The temperature was then ramped to 37 *°*C at a heating rate of 3 *°*C min^−1^ and the measurement continues isothermal for 15 min. Finally, frequency-sweep, from 0.01 to 250 rad/s, was performed at a shear strain amplitude of 1 % at 37 *°*C. The sample then remained at 37 *°*C at a 1.0 % strain for 5 min. A step stress relaxation at a constant shear strain amplitude of 10 % was performed, and relaxation was followed for at least 12 h. Each experimental condition was measured in at least triplicates, *n* ≥ 3. In relaxation profile plots, the normalized data is presented as the average and shaded area represents the standard deviation.

### 4.3 Modeling

All modeling was performed through Matlab R2025a. First, all exported files were separated into each of the rheometry steps, and the stress relaxation step was isolated. Each stress relaxation experiment was normalized with respect to initial stress and smoothed through a standard moving average algorithm. Once normalized, the average stress relaxation profile was calculated for each experimental condition for the first 12 h of relaxation. To assess the characteristic relaxation time (*τ*_0_) at *q* = 1 (40 kDa-HA), the normalized averaged relaxation data of this condition was fitted to the Modified Maxwell Model,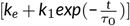, where *k*_*e*_, *k*_1_, and *τ*_0_ were optimized fitting parameters, finding the *τ*_0_ *≈* 28661. Then, all conditions were fitted to the Modified Maxwell Model with a their respective set *τ*_*ϵ*_ = *q · τ*_0_, and *k*_*e*_, *k*_1_ were the optimized fitting parameters. When fitting the Two-Component DCC Network model, 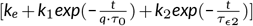, the same set *τ*_*ϵ*_ was maintained, and *k*_*e*_, *k*_1_, *k*_2_, and *τ*_*ϵ*2_ were the optimized fitting parameters. All averaged empirical data was fitted to the corresponding models through a non-linear least-squares local solver, *‘lsqcurvefit’*, which was run 500 times, through a GlobalOptimSolution combined with a MultiStart, to ensure the optimal solution despite potential multiple local minima. As the fitted parameters are representative of physical properties, all fitting parameters (*k*_*e*_, *k*_1_, *k*_2_, and *τ*_*ϵ*2_) were constrained to be ≥ 0.

### 4.4 iPSC-hNPC differentiation and encapsulation

We differentiated human induced pluripotent stem cells (hiPSCs) into neural progenitor cells through a dual SMAD inhibition kit (Stem Cell Technologies).Huang et al. (2025b) After 13 days of differentiation, we dissociated all cells with Accutase (Sigma-Aldrich), and passaged onto plates coated with poly-d-lysine (50 *µ*g/ml; Sigma-Aldrich) and laminin (10 *µ*g/ml; Roche).Huang et al. (2025b) From day 13 onward, the differentiated NPCs were maintained in N3 culture medium, which is composed of 1:1 DMEM/F12 (Thermo Fisher):Neurobasal (Thermo Fisher), N2 supplement (1%, Thermo Fisher), B27 supplement (2%, Thermo Fisher), GlutaMAX (1%, Thermo Fisher), MEM non-essential amino acids (1%, Thermo Fisher), and Penicillin–Streptomycin (1%, Thermo Fisher).Huang et al. (2025b) All NPCs were encapsulated on days 18 - 22 of differentiation. Before encapsulation, the cells were dissociated with Accutase, pelleted by centrifugation, and counted. The NPC pellets were resuspended in 2 wt% ELP-HYD solution at two times the selected final cell density within the gel (i.e., 60,000 cells/*µ*l for a final cell density of 30,000 cells/*µ*l). The cell suspensions were thoroughly mixed with the selected 2 wt% HA-BZA solution, and 10 *µ*l gels were cast in 4 mm diameter, 0.8 mm height silicone molds. The molds were placed into 24-well plates.Huang et al. (2025b) The gels crosslinked for 5 minutes on ice, 15 minutes at room temperature (RT), and 10 minutes at 37 *°*C. Then, we added the N3 media supplemented with 10 µM Y27632 (Tocris Biosciences).Huang et al. (2025b) Fresh media without Y276332 was replenished the next day. Media changes were performed daily for 7 days.

### 4.5 Cell viability

At day 7 (end-point), Live/Dead kit (Invitrogen) was used to assess cell viability at 37 *°*C. Encapsulated gels were washed in phosphate buffered saline (PBS, Invitrogen) for 5 min per wash for three washes total, and incubated with 4.0 *µ*M calcein- AM and 4.0 *µ*M ethidium homodimer- 1 in PBS for 30 min.Navarro et al. (2022) All samples were rinsed in PBS, and imaged on a confocal microscope (Leica STELLARIS 5) at 10x and 20x magnification. Z-stacks of 100 - 150 *µ*m height were taken with 10- *µ*m z-steps at three different locations in each hydrogel. Cell viability was quantified using Fiji software (NIH, version 2.14) by applying intensity thresholds to differentiate and count live cells (green) from dead cells (red).Navarro et al. (2022)

### 4.6 Immunocytochemistry

At the end-point, the gel-encapsulated cells were fixed in 4% paraformaldehyde in PBS for 20 min at RT. The samples were then washed three times with PBS and stored at 4 *°*C until use. The fixed samples were permeabilized with 0.25% (v/v) Triton X-100 in PBS (PBS-T) for 1 hour at RT and then blocked with 5% (w/v) bovine serum albumin (BSA; Roche), 5% (v/v) goat serum (Gibco), and 0.5% Triton X-100 in PBS for 3 hours at RT. Huang et al. (2025b) An antibody solution was prepared in PBS with 2.5% (w/v) BSA, 2.5% (v/v) goat serum, and 0.5% (v/v) Triton X-100. The following primary antibodies were used accross all conditions: mouse anti-*β*3-tubulin (1:400, Cell Signaling, mAb 4466) and rabbit anti-Sox2 (1:400, Cell Signaling, mAb 23064). Primary antibody incubation was performed for 48h at 4 *°*C. After two nights, all samples were washed three times with PBS-T for 30 min each, and then secondary antibodies were diluted in antibody dilution solution as follows: goat anti-rabbit Alexa Fluor 488 (1:500, Invitrogen, A-11008), goat anti-mouse Alexa Fluor 555 (1:500, Invitrogen, A-21422). 4′,6-diamidino-2-phenylindole (DAPI; 1 *µ*g/mL, Cell Signaling, 4083s) was included to stain for nuclei. Huang et al. (2025b) Secondary antibodies were incubated overnight at 4 *°*C. The following day, samples were washed three times with PBS-T for 20 min each and then mounted onto 1 coverslips with ProLong Gold AntiFade Mountant (Thermo Fisher). Huang et al. (2025b) After curing for 24 hours, stained hydrogels were imaged with a confocal microscope (Leica STELLARIS 5, Las-X software). All images were taken at a z-depth of at least 50 *µ*m from the coverslips to avoid possible confounds imparted by the mechanical properties of the glass. Z-stacks of 250 *µ*m height were taken with 10-*µ*m z-steps at five different locations in each hydrogel. Huang et al. (2025b)

### 4.2 Image Analysis

The length of individual neurites was characterized by manually tracing TUBB3-expressing projections, and quantified with the SNT toolbox Arshadi et al. (2021). Cluster size was quantified using CellProfiler based on DAPI staining, identifying objects > 40 pixel units in diameter using no method to distinguish between clumped objects. CellProfiler was also used to quantified the number of Sox2+ cells and area occupied by TUBB3+ neurites. Huang et al. (2025b) Briefly, nuclei and cells were immunolabeled with the relevant markers or antibodies and identified using”IdentifyPrimaryObjects” with the”Minimum Cross-Entropy” thresholding method. TUBB3-labeled neurites were identified by passing the TUBB3 images through “EnhanceOrSuppressFeatures”, “IdentifyPrimaryObjects”, and “MeasureImageAreaOccupied”.Huang et al. (2025b) All quantification on a per cell basis was done by normalizing either the Sox2 count or the area of TUBB3 expression by nuclei count.

### 4.8 Hydrogel erosion

For the erosion study, gels with a volume of 40 *µ*L were cast in the bottom of 0.5 mL Eppendorf tubes following the protocol described above. After gelation, 200 *µ*L of PBS was added on top of the gels and the Eppendorfs were closed, and incubated at 37 *°*C. The entire volume of supernatant was removed and replaced at each experimental timepoint and the supernatant stored –80 ^*°*^C prior to analysis. A uronic assay was used to quantify the amount of HA released into the supernatant, as described earlier Cesaretti et al. (2003). Unmodified HA samples of the same *M*_*w*_ were used as standards and PBS-filled wells used to blank the readings.

## ACKNOWLEDGMENTS

The authors thank Pedro Pinheiro for assistance with the code implementation, Prof. Gerald G. Fuller for helpful discussion, and Dr. Patrik K. Johansson for assistance with image aquisition. This work was supported by the National Science Foundation (NSF) grants DGE-1656518 (N.J.B. and M.S.H.) and DMR-2427971 (S.C.H.); the National Institutes of Health (NIH) grants F31-HL175888 (N.J.B.), F31-NS132505 (M.S.H.), and R01-EY035697, R01-HL173056, R01-MH137333 (S.C.H.); Stanford Cardiovascular Institute seed grant (S.C.H.); the ARCS Foundation Scholarship (N.J.B.); the PhRMA Foundation Predoctoral Fellowship in Drug Delivery (N.J.B.); the American Heart Association (AHA) Predoctoral Fellowship (24PRE1191604) (N.d.P.N.); the Gerald J. Lieberman Fellowship (N.d.P.N, and M.S.H.); the Swiss National Science Foundation Postdoc Mobility Fellowship (PN210723) (F.C.); the Sarafan ChEM-H Chemistry-Biology Interface Program (M.S.H.); and Fundación Alfonso Martin Escudero Fellowship (C.H.L.).

## SUPPORTING INFORMATION

Additional supporting information may be found in the online version of the article at the publisher’s website.

## REFERENCES

Abuwatfa, W.H., Pitt, W.G. & Husseini, G.A. (2024) Scaffold-based 3d cell culture models in cancer research.

Anseth, K.S. & Klok, H.A. (2016) Click chemistry in biomaterials, nanomedicine, and drug delivery. Biomacromolecules, 17(1), 1–3. doi:10.1021/acs.biomac.5b01660, anseth, Kristi S Klok, Harm-Anton eng Editorial 2016/01/12 Biomacromolecules. 2016 Jan 11;17(1):1–3. doi: 10.1021/acs.biomac.5b01660. URL https://www.ncbi.nlm.nih.gov/pubmed/26750314

Arshadi, C., Günther, U., Eddison, M., Harrington, K.I. & Ferreira, T.A. (2021) Snt: a unifying toolbox for quantification of neuronal anatomy. Nature Methods, 18, 374–377. doi:10.1038/s41592-021-01105-7.

Borelli, A.N., Young, M.W., Kirkpatrick, B.E., Jaeschke, M.W., Mellett, S., Porter, S. et al. (2022) Stress relaxation and composition of hydrazone-crosslinked hybrid biopolymer-synthetic hydrogels determine spreading and secretory properties of mscs. Advanced Healthcare Materials, 11(14), 2200393. doi:10.1002/adhm.202200393. URL https://advanced.onlinelibrary.wiley.com/doi/abs/10.1002/adhm.202200393

Cai, P.C., Krajina, B.A. & Spakowitz, A.J. (2020) Brachiation of a polymer chain in the presence of a dynamic network. Phys Rev E, 102(2-1), 020501. doi:10.1103/PhysRevE.102.020501, cai, Pamela C Krajina, Brad A Spakowitz, Andrew J eng 2020/09/19 Phys Rev E. 2020 Aug;102(2-1):020501. doi: 10.1103/PhysRevE.102.020501. URL https://www.ncbi.nlm.nih.gov/pubmed/32942387

Cai, P.C., Su, B., Zou, L., Webber, M.J., Heilshorn, S.C. & Spakowitz, A.J. (2022) Rheological characterization and theoretical modeling establish molecular design rules for tailored dynamically associating polymers. ACS Central Science, 8, 1318–1327. doi:10.1021/acscentsci.2c00432.

Calhoun, M.A., Bentil, S.A., Elliott, E., Otero, J.J., Winter, J.O. & Dupaix, R.B. (2019) Beyond linear elastic modulus: viscoelastic models for brain and brain mimetic hydrogels. ACS Biomaterials Science & Engineering, 5(8), 3964–3973.

Cesaretti, M., Luppi, E., Maccari, F. & Volpi, N. (2003) A 96-well assay for uronic acid carbazole reaction. Carbohydrate Polymers, 54, 59–61. doi:10.1016/S0144-8617(03)00144-9.

Chaudhuri, O. (2017) Viscoelastic hydrogels for 3d cell culture. Biomaterials science, 5(8), 1480–1490.

Chaudhuri, O., Cooper-White, J., Janmey, P.A., Mooney, D.J. & Shenoy, V.B. (2020) Effects of extracellular matrix viscoelasticity on cellular behaviour.

Chaudhuri, O., Gu, L., Darnell, M., Klumpers, D., Bencherif, S.A., Weaver, J.C. et al. (2015) Substrate stress relaxation regulates cell spreading. Nature Communications, 6. doi:10.1038/ncomms7365.

Chaudhuri, O., Gu, L., Klumpers, D., Darnell, M., Bencherif, S.A., Weaver, J.C. et al. (2016) Hydrogels with tunable stress relaxation regulate stem cell fate and activity. Nature Materials, 15, 326–334. doi:10.1038/nmat4489.

Cheng, N., Zhang, Y., Wu, Y., Li, B., Wang, H., Chen, S. et al. (2022) Hydrogel platform capable of molecularly resolved pulling on cells for mechanotransduction. Mater Today Bio, 17, 100476. doi:10.1016/j.mtbio.2022.100476, cheng, Nan Zhang, Yile Wu, Yukai Li, Bohan Wang, Hong Chen, Shaojie Zhao, Peng Cui, Jiaxi Shen, Xiaoqin Zhu, Xingjun Zheng, Yijun eng England 2022/11/22 Mater To-day Bio. 2022 Nov 4;17:100476. doi: 10.1016/j.mtbio.2022.100476. eCollection 2022 Dec 15. URL https://www.ncbi.nlm.nih.gov/pubmed/36407911

Chesselet, M.F. & Carmichael, S.T. (2012) Animal models of neurological disorders. Neurotherapeutics, 9(2), 241–244.

Courbot, O. & Elosegui-Artola, A. (2025) The role of extracellular matrix viscoelasticity in development and disease. npj Biological Physics and Mechanics, 2. doi:10.1038/s44341-025-00014-6.

Dai, X., Wu, D., Xu, K., Ming, P., Cao, S. & Yu, L. (2025) Viscoelastic mechanics: From pathology and cell fate to tissue regeneration biomaterial development.

Darnell, M., Young, S., Gu, L., Shah, N., Lippens, E., Weaver, J. et al. (2017) Substrate stress-relaxation regulates scaffold remodeling and bone formation in vivo. Advanced Healthcare Materials, 6. doi:10.1002/adhm.201601185.

de Gennes, P.G. (1971) Reptation of a polymer chain in the presence of fixed obstacles. The Journal of Chemical Physics, 55(2), 572–579. doi:10.1063/1.1675789. URL https://doi.org/10.1063/1.1675789

de Paiva Narciso, N., Navarro, R.S., Gilchrist, A.E., Trigo, M.L., Rodriguez, G.A. & Heilshorn, S.C. (2023) Design parameters for injectable biopolymeric hydrogels with dynamic covalent chemistry crosslinks. Advanced Healthcare Materials, 12. doi:10.1002/adhm.202301265.

Eliot, L. & Richardson, S.S. (2016) Sex in context: limitations of animal studies for addressing human sex/gender neurobehavioral health disparities. Journal of Neuroscience, 36(47), 11823–11830.

Elosegui-Artola, A., Gupta, A., Najibi, A.J., Seo, B.R., Garry, R., Tringides, C.M. et al. (2023) Matrix viscoelasticity controls spatiotemporal tissue organization. Nature Materials, 22, 117–127. doi:10.1038/s41563-022-01400-4.

Findley, W.N. & Davis, F.A. (2013) Creep and relaxation of nonlinear viscoelastic materials. : Courier corporation.

Hachet, E., Van Den Berghe, H., Bayma, E., Block, M.R. & Auzely-Velty, R. (2012) Design of biomimetic cell-interactive substrates using hyaluronic acid hydrogels with tunable mechanical properties. Biomacromolecules, 13(6), 1818–27. doi:10.1021/bm300324m, hachet, Emilie Van Den Berghe, Helene Bayma, Eric Block, Marc R Auzely-Velty, Rachel eng Research Support, Non-U.S. Gov’t 2012/05/09 Biomacromolecules. 2012 Jun 11;13(6):1818–27. doi: 10.1021/bm300324m. Epub 2012 May 17. URL https://www.ncbi.nlm.nih.gov/pubmed/22559074

Hafeez, S., Ooi, H.W., Morgan, F.L.C., Mota, C., Dettin, M., Van Blitterswijk, C. et al. (2018) Viscoelastic oxidized alginates with reversible imine type crosslinks: Self-healing, injectable, and bioprintable hydrogels. Gels, 4(4). doi:10.3390/gels4040085, hafeez, Shahzad Ooi, Huey Wen Morgan, Francis L C Mota, Carlos Dettin, Monica Van Blitterswijk, Clemens Moroni, Lorenzo Baker, Matthew B eng 694801/H2020 European Research Council/731.016.202/Nederlandse Organisatie voor Wetenschappelijk Onderzoek/Switzerland 2019/01/25 Gels. 2018 Nov 21;4(4):85. doi: 10.3390/gels4040085. URL https://www.ncbi.nlm.nih.gov/pubmed/30674861

Han, Y., Cao, Y. & Lei, H. (2022) Dynamic covalent hydrogels: Strong yet dynamic. Gels, 8(9). doi:10.3390/gels8090577, han, Yueying Cao, Yi Lei, Hai eng T2222019, 11974174, 11934008/National Natural Science Foundation of China/2020YFA0908100/National Key Ramp;D Program of China/No number/State Key Laboratory of Precision Measurement Technology and Instruments (Tianjin University)/Review Switzerland 2022/09/23 Gels. 2022 Sep 10;8(9):577. doi: 10.3390/gels8090577. URL https://www.ncbi.nlm.nih.gov/pubmed/36135289

Hefferon, M.E., Huang, M.S., Liu, Y., Navarro, R.S., de Paiva Narciso, N., Zhang, D. et al. (2023) Cell microencapsulation within engineered hyaluronan elastin-like protein (help) hydrogels. Current Protocols, 3. doi:10.1002/cpz1.917.

Huang, D., Huang, Y., Xiao, Y., Yang, X., Lin, H., Feng, G. et al. (2019) Viscoelasticity in natural tissues and engineered scaffolds for tissue reconstruction.

Huang, M.S., Christakopoulos, F., Roth, J.G. & Heilshorn, S.C. (2025) Organoid bioprinting: from cells to functional tissues. Nature Reviews Bioengineering, 3, 126–142. doi:10.1038/s44222-024-00268-0. URL https://www.nature.com/articles/s44222-024-00268-0

Huang, M.S., LeSavage, B.L., Ghorbani, S., Gilchrist, A.E., Roth, J.G., Huerta-López, C. et al. (2025) Viscoelastic n-cadherin-like interactions maintain neural progenitor cell stemness within 3d matrices. Nature Communications, 16. doi:10.1038/s41467-025-60540-8.

Huang, Y., Liu, T., Huang, Q. & Wang, Y. (2024) From organ-on-a-chip to human-on-a-chip: A review of research progress and latest applications.

Hull, S.M., Lou, J., Lindsay, C.D., Navarro, R.S., Cai, B., Brunel, L.G. et al. (2023) 3d bioprinting of dynamic hydrogel bioinks enabled by small molecule modulators. Sci Adv, 9(13), eade7880. doi:10.1126/sciadv.ade7880, hull, Sarah M Lou, Junzhe Lindsay, Christopher D Navarro, Renato S Cai, Betty Brunel, Lucia G Westerfield, Ashley D Xia, Yan Heilshorn, Sarah C eng 2023/04/01 Sci Adv. 2023 Mar 31;9(13):eade7880. doi: 10.1126/sciadv.ade7880. Epub 2023 Mar 31. URL https://www.ncbi.nlm.nih.gov/pubmed/37000873

Hunt, D.R., Klett, K.C., Mascharak, S., Wang, H., Gong, D., Lou, J. et al. (2021) Engineered matrices enable the culture of human patient-derived intestinal organoids. Adv Sci (Weinh), 8(10), 2004705. doi:10.1002/advs.202004705, hunt, Daniel R Klett, Katarina C Mascharak, Shamik Wang, Huiyuan Gong, Diana Lou, Junzhe Li, Xingnan Cai, Pamela C Suhar, Riley A Co, Julia Y LeSavage, Bauer L Foster, Abbygail A Guan, Yuan Amieva, Manuel R Peltz, Gary Xia, Yan Kuo, Calvin J Heilshorn, Sarah C eng U19 AI116484/AI/NIAID NIH HHS/R01 DK115728/DK/NIDDK NIH HHS/R01 DK102182/DK/NIDDK NIH HHS/U01 DA044399/DA/NIDA NIH HHS/R21 HL138042/HL/NHLBI NIH HHS/R01 HL142718/HL/NHLBI NIH HHS/R01 EB027171/EB/NIBIB NIH HHS/T32 GM119995/GM/NIGMS NIH HHS/Research Support, N.I.H., Extramural Research Support, U.S. Gov’t, Non-P.H.S. Germany 2021/05/25 Adv Sci (Weinh). 2021 Mar 12;8(10):2004705. doi: 10.1002/advs.202004705. eCollection 2021 May. URL https://www.ncbi.nlm.nih.gov/pubmed/34026461

Ingber, D.E. (2022) Human organs-on-chips for disease modelling, drug development and personalized medicine.

Jaeschke, M.W., Borelli, A.N., Skillin, N.P., White, T.J. & Anseth, K.S. (2024) Engineering a hydrazone and triazole crosslinked hydrogel for extrusion-based printing and cell delivery. Advanced Healthcare Materials, 13(20), 2400062. doi:10.1002/adhm.202400062. URL https://advanced.onlinelibrary.wiley.com/doi/abs/10.1002/adhm.202400062

Kalin, N.H. (2021) Understanding the value and limitations of mri neuroimaging in psychiatry.

Kim, J., Koo, B.K. & Knoblich, J.A. (2020) Human organoids: model systems for human biology and medicine.

LeSavage, B.L., Suhar, N.A., Madl, C.M. & Heilshorn, S.C. (2018) Production of elastin-like protein hydrogels for encapsulation and immunostaining of cells in 3d. J Vis Exp, (135), 57739. doi:10.3791/57739, leSavage, Bauer L Suhar, Nicholas A Madl, Christopher M Heilshorn, Sarah C eng F31 EB020502/EB/NIBIB NIH HHS/U19 AI116484/AI/NIAID NIH HHS/T32 GM008412/GM/NIGMS NIH HHS/R21 EB018407/EB/NIBIB NIH HHS/P2C HD086843/HD/NICHD NIH HHS/Research Support, N.I.H., Extramural Research Support, Non-U.S. Gov’t Research Support, U.S. Gov’t, Non-P.H.S. Video-Audio Media 2018/06/05 J Vis Exp. 2018 May 19;(135):57739. doi: 10.3791/57739. URL https://www.ncbi.nlm.nih.gov/pubmed/29863669

LeSavage, B.L., Zhang, D., Huerta-López, C., Gilchrist, A.E., Krajina, B.A., Karlsson, K. et al. (2024) Engineered matrices reveal stiffness-mediated chemoresistance in patient-derived pancreatic cancer organoids. Nature Materials, 23(8), 1138–1149. doi:10.1038/s41563-024-01908-x. URL https://doi.org/10.1038/s41563-024-01908-x

Leung, C.M., de Haan, P., Ronaldson-Bouchard, K., Kim, G.A., Ko, J., Rho, H.S. et al. (2022) A guide to the organ-on-a-chip.

Lin, Y.H., Lou, J., Xia, Y. & Chaudhuri, O. (2024) Cross-linker architectures impact viscoelasticity in dynamic covalent hydrogels. Advanced Healthcare Materials, 13(30), 2402059. doi:10.1002/adhm.202402059. URL https://advanced.onlinelibrary.wiley.com/doi/abs/10.1002/adhm.202402059

Liu, D.D., He, J.Q., Sinha, R., Eastman, A.E., Toland, A.M., Morri, M. et al. (2023) Purification and characterization of human neural stem and progenitor cells. Cell, 186, 1179–1194.e15. doi:10.1016/j.cell.2023.02.017.

Liu, X., Zhou, Z., Zhang, Y., Zhong, H., Cai, X. & Guan, R. (2025) Recent progress on the organoids: Techniques, advantages and applications.

Liu, Y., Gilchrist, A.E. & Heilshorn, S.C. (2024) Engineered protein hydrogels as biomimetic cellular scaffolds. Advanced Materials, 36(45), 2407794. doi:10.1002/adma.202407794. URL https://advanced.onlinelibrary.wiley.com/doi/abs/10.1002/adma.202407794

Liu, Y., Gilchrist, A.E., Johansson, P.K., Guan, Y., Deras, J.D., Liu, Y.C. et al. (2025) Engineered hydrogels for organoid models of human nonalcoholic fatty liver disease. Advanced Science, 12, e17332. doi:10.1002/ADVS.202417332. URL https://pmc.ncbi.nlm.nih.gov/articles/PMC12165117/

Ma, Z., Obuseh, F.O., Freedman, B.R., Kim, J., Torre, M. & Mooney, D.J. (2025) Integrating hydrogels and biomedical plastics via in situ physical entanglements and covalent bonding. Advanced Healthcare Materials, 14(4), 2402605. doi:10.1002/adhm.202402605. URL https://advanced.onlinelibrary.wiley.com/doi/abs/10.1002/adhm.202402605

Madl, C.M. & Heilshorn, S.C. (2018) Bioorthogonal strategies for engineering extracellular matrices. Adv Funct Mater, 28(11). doi:10.1002/adfm.201706046, madl, Christopher M Heilshorn, Sarah C eng R21 EB020235/EB/NIBIB NIH HHS/R21 HL138042/HL/NHLBI NIH HHS/U19 AI116484/AI/NIAID NIH HHS/Germany 2018/03/14 Adv Funct Mater. 2018 Mar 14;28(11):1706046. doi: 10.1002/adfm.201706046. Epub 2018 Jan URL https://www.ncbi.nlm.nih.gov/pubmed/31558890

McKinnon, D.D., Domaille, D.W., Cha, J.N. & Anseth, K.S. (2014) Biophysically defined and cytocompatible covalently adaptable networks as viscoelastic 3d cell culture systems. Adv Mater, 26(6), 865–72. doi:10.1002/adma.201303680, mcKinnon, Daniel D Domaille, Dylan W Cha, Jennifer N Anseth, Kristi S eng HHMI/Howard Hughes Medical Institute/T32 GM008732/GM/NIGMS NIH HHS/5 T32 GM 8732/GM/NIGMS NIH HHS/Research Support, N.I.H., Extramural Research Support, Non-U.S. Gov’t Research Support, U.S. Gov’t, Non-P.H.S. Validation Study Germany 2013/10/16 Adv Mater. 2014 Feb 12;26(6):865–72. doi: 10.1002/adma.201303680. Epub 2013 Oct 11. URL https://www.ncbi.nlm.nih.gov/pubmed/24127293

Muir, V.G. & Burdick, J.A. (2021) Chemically modified biopolymers for the formation of biomedical hydrogels. Chem Rev, 121(18), 10908–10949. doi:10.1021/acs.chemrev.0c00923, muir, Victoria G Burdick, Jason A eng R01 AR056624/AR/NIAMS NIH HHS/R01 AR077362/AR/NIAMS NIH HHS/Research Support, N.I.H., Extramural Research Support, Non-U.S. Gov’t Review 2020/12/29 Chem Rev. 2021 Sep 22;121(18):10908–10949. doi: 10.1021/acs.chemrev.0c00923. Epub 2020 Dec 23. URL https://www.ncbi.nlm.nih.gov/pubmed/33356174

Navarro, R.S., Huang, M.S., Roth, J.G., Hubka, K.M., Long, C.M., Enejder, A. et al. (2022) Tuning polymer hydrophilicity to regulate gel mechanics and encapsulated cell morphology. Adv Healthc Mater, 11(13), e2200011. doi:10.1002/adhm.202200011, navarro, Renato S Huang, Michelle S Roth, Julien G Hubka, Kelsea M Long, Chris M Enejder, Annika Heilshorn, Sarah C eng R01 HL151997/HL/NHLBI NIH HHS/R01 EB027171/EB/NIBIB NIH HHS/R21 NS114549/NS/NINDS NIH HHS/R01 HL142718/HL/NHLBI NIH HHS/Research Support, N.I.H., Extramural Research Support, Non-U.S. Gov’t Research Support, U.S. Gov’t, Non-P.H.S. Germany 2022/04/05 Adv Healthc Mater. 2022 Jul;11(13):e2200011. doi: 10.1002/adhm.202200011. Epub 2022 May 6. URL https://www.ncbi.nlm.nih.gov/pubmed/35373510

Oelschlaeger, C., Cota Pinto Coelho, M. & Willenbacher, N. (2013) Chain flexibility and dynamics of polysaccharide hyaluronan in entangled solutions: a high frequency rheology and diffusing wave spectroscopy study. Biomacromolecules, 14(10), 3689–96. doi:10.1021/bm4010436, oelschlaeger, C Cota Pinto Coelho, M Willenbacher, N eng 2013/08/29 Biomacromolecules. 2013 Oct 14;14(10):3689–96. doi: 10.1021/bm4010436. Epub 2013 Sep 11. URL https://www.ncbi.nlm.nih.gov/pubmed/23980898

Rosa, M.E.D. & Winter, H.H. (1994) The effect of entanglements on the rheological behavior of polybutadiene critical gels. Rheologica Acta, 33, 220–237. doi:10.1007/BF00437307. URL https://doi.org/10.1007/BF00437307

Roth, J.G., Huang, M.S., Navarro, R.S., Akram, J.T., Lesavage, B.L. & Heilshorn, S.C. (2023) Tunable hydrogel viscoelasticity modulates human neural maturation. Science Advances„ eadh8313. URL https://www.science.org

Rubinstein, M., Colby, R.H. & Knovel (2003) Polymer physics. Oxford ; New York: Oxford University Press. (Firm) Includes bibliographical references and index. 1Introduction –2Ideal chains –3Real chains –4Thermodynamics of mixing –5Polymer solutions –6Random branching and gelatin –7Networks and gels –8Unentangled polymer dynamics –9Entangled polymer dynamics. “This is a polymer physics textbook for upper level undergraduates and first year graduate students. Any student with a working knowledge of calculus, physics, and chemistry will be able to read this book. The essential tools of the polymer physical chemist or engineer are derived in this book without skipping any steps. The book is a self-contained treatise that could also serve as a useful reference for scientists and engineers working with polymers. While no prior knowledge of polymers is assumed, the book goes far beyond introductory polymer terms in the scope of what is covered. The fundamental concepts required to fully understand polymer melts, solutions and gels in terms of both static structure and dynamics are explained in detail. Problems at the end of each chapter provide the reader with the opportunity to apply what has been learned to practice.”–BOOK JACKET. Description based on print version record. Restricted to users at subscribing institutions. [electronic resource]. URL http://app.knovel.com/web/toc.v/cid:kpPP000001 http://mirlyn.lib.umich.edu/Record/011740185

Rubinstein, M. & Panyukov, S. (2002) Elasticity of polymer networks. Macromolecules, 35, 6670–6686. doi:10.1021/ma0203849, doi: 10.1021/ma0203849. URL https://doi.org/10.1021/ma0203849

Rubinstein, M. & Semenov, A.N. (2001) Dynamics of entangled solutions of associating polymers. Macromolecules, 34(4), 1058–1068. doi:10.1021/ma0013049, 400ef Times Cited:406 Cited References Count:18. URL <GotoISI>://WOS:000166858700052

Stiles, J. & Jernigan, T.L. (2010) The basics of brain development.

Suhar, R.A., Huang, M.S., Navarro, R.S., Rodriguez, G.A. & Heilshorn, S.C. (2023) A library of elastin-like proteins with tunable matrix ligands for in vitro 3d neural cell culture. Biomacromolecules, 24, 5926–5939. doi:10.1021/acs.biomac.3c00941.

Tang, X.Y., Wu, S., Wang, D., Chu, C., Hong, Y., Tao, M. et al. (2022) Human organoids in basic research and clinical applications.

Terzopoulou, Z., Zamboulis, A., Koumentakou, I., Michailidou, G., Noordam, M.J. & Bikiaris, D.N. (2022) Biocompatible synthetic polymers for tissue engineering purposes. Biomacromolecules, 23, 1841–1863. doi:10.1021/acs.biomac.2c00047.

Tibbitt, M.W. & Anseth, K.S. (2009) Hydrogels as extracellular matrix mimics for 3d cell culture. Biotechnology and Bioengineering, 103, 655–663. doi:10.1002/bit.22361.

Wang, H., Paul, A., Nguyen, D., Enejder, A. & Heilshorn, S.C. (2018) Tunable control of hydrogel microstructure by kinetic competition between self-assembly and crosslinking of elastin-like proteins. ACS Appl Mater Interfaces, 10(26), 21808–21815. doi:10.1021/acsami.8b02461, wang, Huiyuan Paul, Alexandra Nguyen, Duong Enejder, Annika Heilshorn, Sarah C eng 2018/06/06 ACS Appl Mater Interfaces. 2018 Jul 5;10(26):21808–21815. doi: 10.1021/acsami.8b02461. Epub 2018 Jun 20. URL https://www.ncbi.nlm.nih.gov/pubmed/29869869

Xu, F., Dawson, C., Lamb, M., Mueller, E., Stefanek, E., Akbari, M. et al. (2022) Hydrogels for tissue engineering: Addressing key design needs toward clinical translation. Frontiers in Bioengineering and Biotechnology, 10. doi:10.3389/fbioe.2022.849831.

Xu, X., Jha, A.K., Harrington, D.A., Farach-Carson, M.C. & Jia, X. (2012) Hyaluronic acid-based hydrogels: from a natural polysaccharide to complex networks. Soft Matter, 8(12), 3280–3294. doi:10.1039/C2SM06463D, xu, Xian Jha, Amit K Harrington, Daniel A Farach-Carson, Mary C Jia, Xinqiao eng P01 CA098912/CA/NCI NIH HHS/P20 RR017716/RR/NCRR NIH HHS/R01 DC008965/DC/NIDCD NIH HHS/R01 DC008965-05/DC/NIDCD NIH HHS/England 2012/03/16 Soft Matter. 2012;8(12):3280–3294. doi: 10.1039/C2SM06463D. URL https://www.ncbi.nlm.nih.gov/pubmed/22419946

Zhang, V., Accardo, J.V., Kevlishvili, I., Woods, E.F., Chapman, S.J., Eckdahl, C.T. et al. (2023) Tailoring dynamic hydrogels by controlling associative exchange rates. Chem, 9, 2298–2317. doi:10.1016/j.chempr.2023.05.018.

