## Supplemental Information for "Tuning viscoelasticity of dynamic covalent hydrogels for human tissue modeling"

**List of Supplemental Information:**

- **Figure S1.**Representative NMR for Hyaluronic Acid (60 kDa) at different degrees of benzaldehyde functionalization (6, 12 and 20%).
- **Table S1.** P-values for storage moduli with different ALD:BZA ratios in Figure 1E.
- **Table S2.**. P-values for storage moduli with different HA Mw in Figure 2C.
- **Table S3.** P-values for fraction of non-dissipated stress after 12 hours with different HA Mw in Figure 2D
- **Table S4.** P-values for time necessary to dissipate 15% of applied stress with different HA Mw in Figure 2E
- **Table S5.** Fitted values for Modified Maxwell Model for HELP formulations in Figure 3E.
- **Table S6.** P-values for time necessary to dissipate 15% of applied stress normalized with respect to q factor for different HA Mw in Figure 3H
- **Table S7.** P-values for storage moduli of 20- and 60-kDa HA Mw HELP formulations in Figure 4E
- **Table S8.** P-values for storage moduli of 500- and 1000-kDa HA Mw HELP formulations in Figure 5A
- **Table S9.** Calculated theoretical entanglement concentrations ($C_e$) for non-functionalized hyaluronic acid
- **Figure S2.** Natural log of fraction of non-dissipated stress ( / 0) against time for formulations with HA Mw 20-, 40-, 60- and 100-kDa (average of n=3-4), compared to the theoretical fit of Two-Polymer DCC Network Model.
- **Table S10.** Fitted values for Two-Polymer DCC Network Model for HELP

formulations in Figure 5G

– **Figure S3.** Comparison of fitted values for Two-Polymer DCC Network Model and Modified Maxwell Model
– **Figure S4.** Live and Dead staining of hiPSC-derived NPCs within selected HELP formulations for 1 week

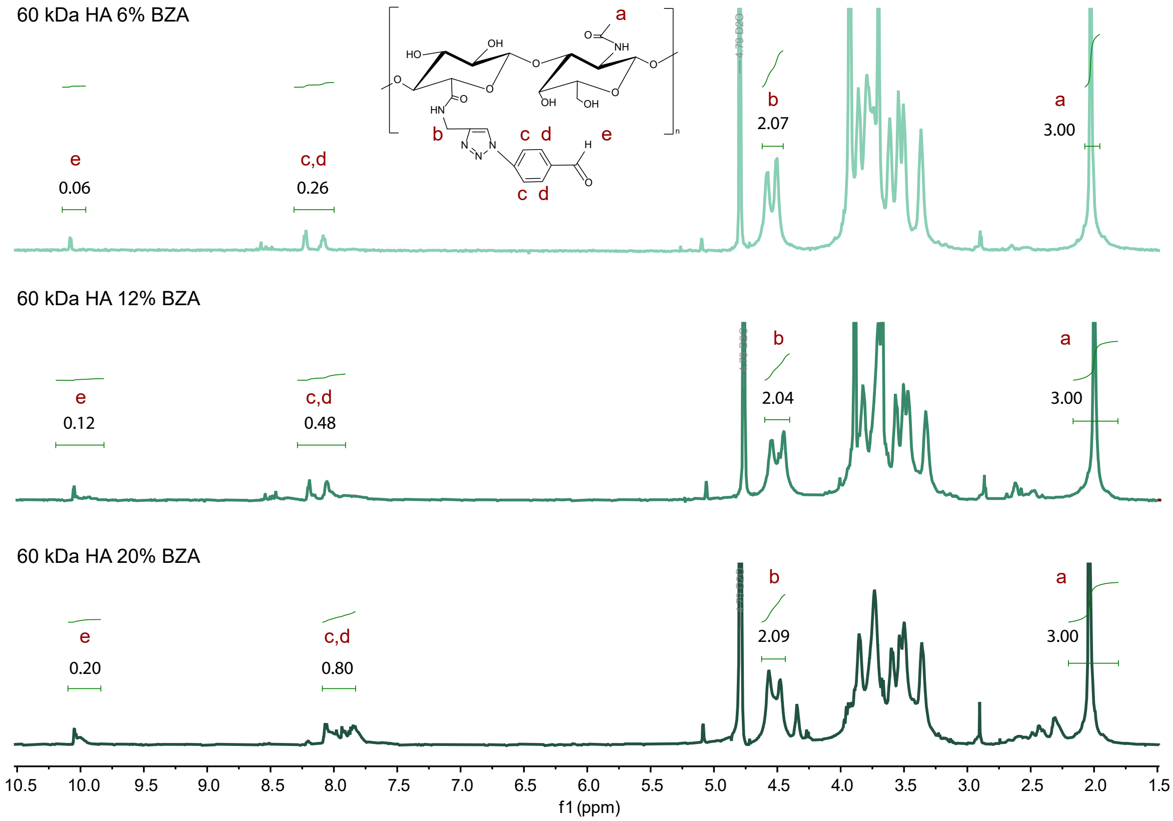


Figure S1: Representative NMR for Hyaluronic Acid (60 kDa) at different degrees of benzaldehyde functionalization (6, 12 and 20%)

Table S1: P-values for storage moduli with different ALD:BZA ratios in Figure 1E.

| Tukey's multiple comparisons test | Summary | Adjusted P Value |
|---|---|---|
| HELP 12% 0:100 vs. HELP 12% 50:50 | ns | 0.8439 |
| HELP 12% 0:100 vs. HELP 12% 100:0 | ns | 0.079 |
| HELP 12% 0:100 vs. HELP 6% 0:100 | **** | <0.0001 |
| HELP 12% 0:100 vs. HELP 6% 50:50 | **** | <0.0001 |
| HELP 12% 0:100 vs. HELP 6% 100:0 | **** | <0.0001 |
| HELP 12% 50:50 vs. HELP 12% 100:0 | ns | 0.4376 |
| HELP 12% 50:50 vs. HELP 6% 0:100 | **** | <0.0001 |
| HELP 12% 50:50 vs. HELP 6% 50:50 | **** | <0.0001 |
| HELP 12% 50:50 vs. HELP 6% 100:0 | **** | <0.0001 |
| HELP 12% 100:0 vs. HELP 6% 0:100 | **** | <0.0001 |
| HELP 12% 100:0 vs. HELP 6% 50:50 | *** | 0.0007 |
| HELP 12% 100:0 vs. HELP 6% 100:0 | **** | <0.0001 |
| HELP 6% 0:100 vs. HELP 6% 50:50 | ns | 0.6581 |
| HELP 6% 0:100 vs. HELP 6% 100:0 | ns | 0.2941 |
| HELP 6% 50:50 vs. HELP 6% 100:0 | * | 0.0184 |

Table S2: P-values for storage moduli with different HA $M_w$ in Figure 2C.

| Tukey's multiple comparisons test | Summary | Adjusted P Value |
|---|---|---|
| HA 60 kDa 12%BZA vs. HA 40 kDa 12%BZA | * | 0.01 |
| HA 60 kDa 12%BZA vs. HA 20 kDa 12%BZA | *** | 0.0002 |
| HA 60 kDa 12%BZA vs. HA 500 kDa 12%BZA | ns | 0.9731 |
| HA 60 kDa 12%BZA vs. HA 100 kDa 12%BZA | * | 0.0429 |
| HA 40 kDa 12%BZA vs. HA 20 kDa 12%BZA | ns | 0.1462 |
| HA 40 kDa 12%BZA vs. HA 500 kDa 12%BZA | ** | 0.0059 |
| HA 40 kDa 12%BZA vs. HA 100 kDa 12%BZA | **** | <0.0001 |
| HA 20 kDa 12%BZA vs. HA 500 kDa 12%BZA | *** | 0.0002 |
| HA 20 kDa 12%BZA vs. HA 100 kDa 12%BZA | **** | <0.0001 |
| HA 500 kDa 12%BZA vs. HA 100 kDa 12%BZA | ns | 0.1767 |

Table S3: P-values for fraction of non-dissipated stress after 12 hours with different HA $M_w$ in Figure 2D

| Tukey's multiple comparisons test | Summary | Adjusted P Value |
|---|---|---|
| HA 20 kDa 12%BZA vs. HA 40 kDa 12%BZA | ns | 0.2737 |
| HA 20 kDa 12%BZA vs. HA 60 kDa 12%BZA | ** | 0.0079 |
| HA 20 kDa 12%BZA vs. HA 100 kDa 12%BZA | * | 0.0191 |
| HA 20 kDa 12%BZA vs. HA 500 kDa 12%BZA | ns | 0.9098 |
| HA 40 kDa 12%BZA vs. HA 60 kDa 12%BZA | ns | 0.2259 |
| HA 40 kDa 12%BZA vs. HA 100 kDa 12%BZA | ns | 0.4627 |
| HA 40 kDa 12%BZA vs. HA 500 kDa 12%BZA | ns | 0.7522 |
| HA 60 kDa 12%BZA vs. HA 100 kDa 12%BZA | ns | 0.9824 |
| HA 60 kDa 12%BZA vs. HA 500 kDa 12%BZA | * | 0.0403 |
| HA 100 kDa 12%BZA vs. HA 500 kDa 12%BZA | ns | 0.0947 |

Table S4: P-values for time necessary to dissipate 15% of applied stress with different HA $M_w$ in Figure 2E

| Tukey's multiple comparisons test | Summary | Adjusted P Value |
|---|---|---|
| HA 20 kDa 12%BZA vs. HA 40 kDa 12%BZA | ns | 0.7576 |
| HA 20 kDa 12%BZA vs. HA 60 kDa 12%BZA | ** | 0.0058 |
| HA 20 kDa 12%BZA vs. HA 100 kDa 12%BZA | ns | 0.1399 |
| HA 20 kDa 12%BZA vs. HA 500 kDa 12%BZA | ns | 0.9996 |
| HA 40 kDa 12%BZA vs. HA 60 kDa 12%BZA | * | 0.0308 |
| HA 40 kDa 12%BZA vs. HA 100 kDa 12%BZA | ns | 0.6006 |
| HA 40 kDa 12%BZA vs. HA 500 kDa 12%BZA | ns | 0.8631 |
| HA 60 kDa 12%BZA vs. HA 100 kDa 12%BZA | ns | 0.3377 |
| HA 60 kDa 12%BZA vs. HA 500 kDa 12%BZA | ** | 0.0085 |
| HA 100 kDa 12%BZA vs. HA 500 kDa 12%BZA | ns | 0.1972 |

Table S5: Fitted values for Modified Maxwell Model for HELP formulations in Figure 3E.

| HA $M_w$ [kDa] | % BZA | $q$ | $\tau_\varepsilon = q \cdot \tau_0$ $[\cdot 10^4]$ | $k_e$ | $k_1$ | $R^2$ |
|---|---|---|---|---|---|---|
| 20 | 12 | 0.5 | 1.43 | 7.27 | 2.62 | 0.989 |
| 20 | 20 | 0.5 | 1.43 | 8.28 | 1.56 | 0.983 |
| 40 | 12 | 1 | 2.87 | 7.08 | 2.89 | 0.999 |
| 60 | 6 | 1.5 | 4.30 | 0.81 | 8.82 | 0.988 |
| 60 | 12 | 1.5 | 4.30 | 7.61 | 2.35 | 0.998 |
| 60 | 20 | 1.5 | 4.30 | 7.95 | 2.09 | 0.981 |
| 100 | 12 | 2.5 | 7.17 | 6.21 | 3.67 | 0.989 |
| 500 | 6 | 12.5 | 35.82 | $1.29 \cdot 10^{-13}$ | 7.46 | 0.152 |
| 500 | 12 | 12.5 | 35.82 | $3.04 \cdot 10^{-14}$ | 9.51 | 0.541 |
| 500 | 20 | 12.5 | 35.82 | $2.37 \cdot 10^{-14}$ | 9.58 | 0.535 |
| 1000 | 12 | 25 | 71.65 | $3.64 \cdot 10^{-14}$ | 9.14 | 0.531 |

Table S6: P-values for time necessary to dissipate 15% of applied stress normalized with respect to $q$ factor for different HA $M_w$ in Figure 3H

| Tukey's multiple comparisons test | Summary | Adjusted P Value |
|---|---|---|
| HA 20 kDa 12%BZA vs. HA 40 kDa 12%BZA | ns | >0.9999 |
| HA 20 kDa 12%BZA vs. HA 60 kDa 12%BZA | ns | 0.2374 |
| HA 20 kDa 12%BZA vs. HA 100 kDa 12%BZA | ns | 0.641 |
| HA 20 kDa 12%BZA vs. HA 500 kDa 12%BZA | * | 0.019 |
| HA 20 kDa 12%BZA vs. HA 1000 kDa 12%BZA | ** | 0.0097 |
| HA 40 kDa 12%BZA vs. HA 60 kDa 12%BZA | ns | 0.1779 |
| HA 40 kDa 12%BZA vs. HA 100 kDa 12%BZA | ns | 0.5633 |
| HA 40 kDa 12%BZA vs. HA 500 kDa 12%BZA | * | 0.0113 |
| HA 40 kDa 12%BZA vs. HA 1000 kDa 12%BZA | ** | 0.005 |
| HA 60 kDa 12%BZA vs. HA 100 kDa 12%BZA | ** | 0.0075 |
| HA 60 kDa 12%BZA vs. HA 500 kDa 12%BZA | *** | 0.0001 |
| HA 60 kDa 12%BZA vs. HA 1000 kDa 12%BZA | **** | <0.0001 |
| HA 100 kDa 12%BZA vs. HA 500 kDa 12%BZA | ns | 0.2032 |
| HA 100 kDa 12%BZA vs. HA 1000 kDa 12%BZA | ns | 0.1257 |
| HA 500 kDa 12%BZA vs. HA 1000 kDa 12%BZA | ns | >0.9999 |

Table S7: P-values for storage moduli of 20- and 60-kDa HA $M_w$ HELP formulations in Figure 4E

| Tukey's multiple comparisons test | Summary | Adjusted P Value |
|---|---|---|
| HA 60 kDa 12%BZA vs. HA 20 kDa 12%BZA | **** | <0.0001 |
| HA 60 kDa 12%BZA vs. HA 60 kDa 6%BZA | **** | <0.0001 |
| HA 60 kDa 12%BZA vs. HA 60 kDa 20%BZA | * | 0.0306 |
| HA 60 kDa 12%BZA vs. HA 20 kDa 20%BZA | **** | <0.0001 |
| HA 20 kDa 12%BZA vs. HA 60 kDa 6%BZA | ns | 0.9879 |
| HA 20 kDa 12%BZA vs. HA 60 kDa 20%BZA | **** | <0.0001 |
| HA 20 kDa 12%BZA vs. HA 20 kDa 20%BZA | ns | 0.9933 |
| HA 60 kDa 6%BZA vs. HA 60 kDa 20%BZA | **** | <0.0001 |
| HA 60 kDa 6%BZA vs. HA 20 kDa 20%BZA | ns | 0.8964 |
| HA 60 kDa 20%BZA vs. HA 20 kDa 20%BZA | **** | <0.0001 |

Table S8: P-values for storage moduli of 500- and 1000-kDa HA $M_w$ HELP formulations in Figure 5A

| Tukey's multiple comparisons test | Summary | Adjusted P Value |
|---|---|---|
| HA 500 kDa 12%BZA vs. HA 500 kDa 20%BZA | *** | 0.0002 |
| HA 500 kDa 12%BZA vs. HA 1 MDa 12%BZA | **** | <0.0001 |
| HA 500 kDa 12%BZA vs. HA 500 kDa 6%BZA | **** | <0.0001 |
| HA 500 kDa 20%BZA vs. HA 1 MDa 12%BZA | ns | 0.2603 |
| HA 500 kDa 20%BZA vs. HA 500 kDa 6%BZA | ** | 0.0012 |
| HA 1 MDa 12%BZA vs. HA 500 kDa 6%BZA | * | 0.0102 |

Table S9: Calculated theoretical entanglement concentrations ($C_e$) for non-functionalized hyaluronic acid

| M.W. (kDa) | $C_e$ (wt%) |
|---|---|
| 20 | 26.15 |
| 40 | 15.40 |
| 60 | 11.30 |
| 100 | 7.65 |
| 500 | 2.24 |
| 1000 | 1.31 |
| 1430 | 1 |

**Entanglement concentration calculation**

Entanglement concentrations can be estimated from the following scaling relationship: $C_e \sim (N_e/N)^{(3\nu-1)}$, where $C_e$ is the entanglement concentration, $N_e$ is the number of Kuhn monomers between entanglements, $N$ is the total number of Kuhn monomers, and $\nu$ is a solvent quality parameter [1]. We assume that our system has a good solvent ($\nu = 0.588$). Using a known entangled hyaluronic acid solution as a reference ($C_e = 10\ g/L = 1\ wt\%$ for a hyaluronic acid chain of molecular weight 1430 kDa) [1-3], we estimated the $C_e$ for a range of other hyaluronic acid molecular weights.

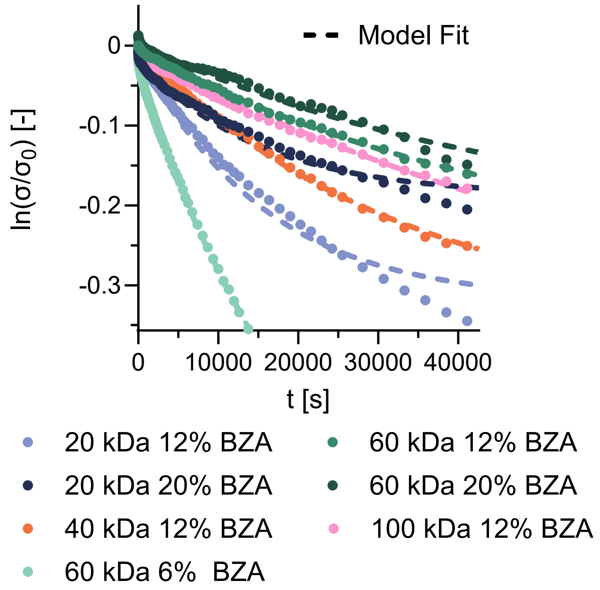


Figure S2: Natural log of fraction of non-dissipated stress (/0) against time for formulations with HA $M_w$ 20-, 40-, 60- and 100-kDa (average of n=3-4), compared to the theoretical fit of Two-Polymer DCC Network Model

Table S10: Fitted values for Two-Polymer DCC Network Model for HELP formulations in Figure 5G

| HA $M_w$ [kDa] | % BZA | $q$ | $\tau_\varepsilon = q \cdot \tau_0$ [$\cdot 10^4$] | $k_e$ | $k_1$ | $k_2$ | $\tau_{\varepsilon 2}$ | $R^2$ |
|---|---|---|---|---|---|---|---|---|
| 20 | 12 | 0.5 | 1.43 | 7.27 | 2.62 | 0.16 | 2.95 | 0.989 |
| 20 | 20 | 0.5 | 1.43 | 8.29 | 1.53 | 0.14 | 58.04 | 0.987 |
| 40 | 12 | 1 | 2.87 | 7.11 | 2.84 | 0.05 | 565.68 | 0.999 |
| 60 | 6 | 1.5 | 4.30 | 1.86 | 7.05 | 0.87 | 4618.92 | 1.000 |
| 60 | 12 | 1.5 | 4.30 | 7.65 | 2.28 | 0.06 | 401.15 | 1.000 |
| 60 | 20 | 1.5 | 4.30 | 8.00 | 2.02 | 0.10 | 156.15 | 0.986 |
| 100 | 12 | 2.5 | 7.17 | 6.40 | 3.43 | 0.13 | 603.12 | 0.999 |
| 500 | 6 | 12.5 | 35.82 | $2.39 \cdot 10^{-13}$ | 1.70 | 7.58 | 13287.92 | 0.996 |
| 500 | 12 | 12.5 | 35.82 | $4.08 \cdot 10^{-13}$ | 8.21 | 1.73 | 12397.92 | 0.999 |
| 500 | 20 | 12.5 | 35.82 | $1.96 \cdot 10^{-13}$ | 7.61 | 2.37 | 22054.97 | 0.997 |
| 1000 | 12 | 25 | 71.65 | $2.57 \cdot 10^{-9}$ | 8.49 | 1.07 | 6548.49 | 0.971 |

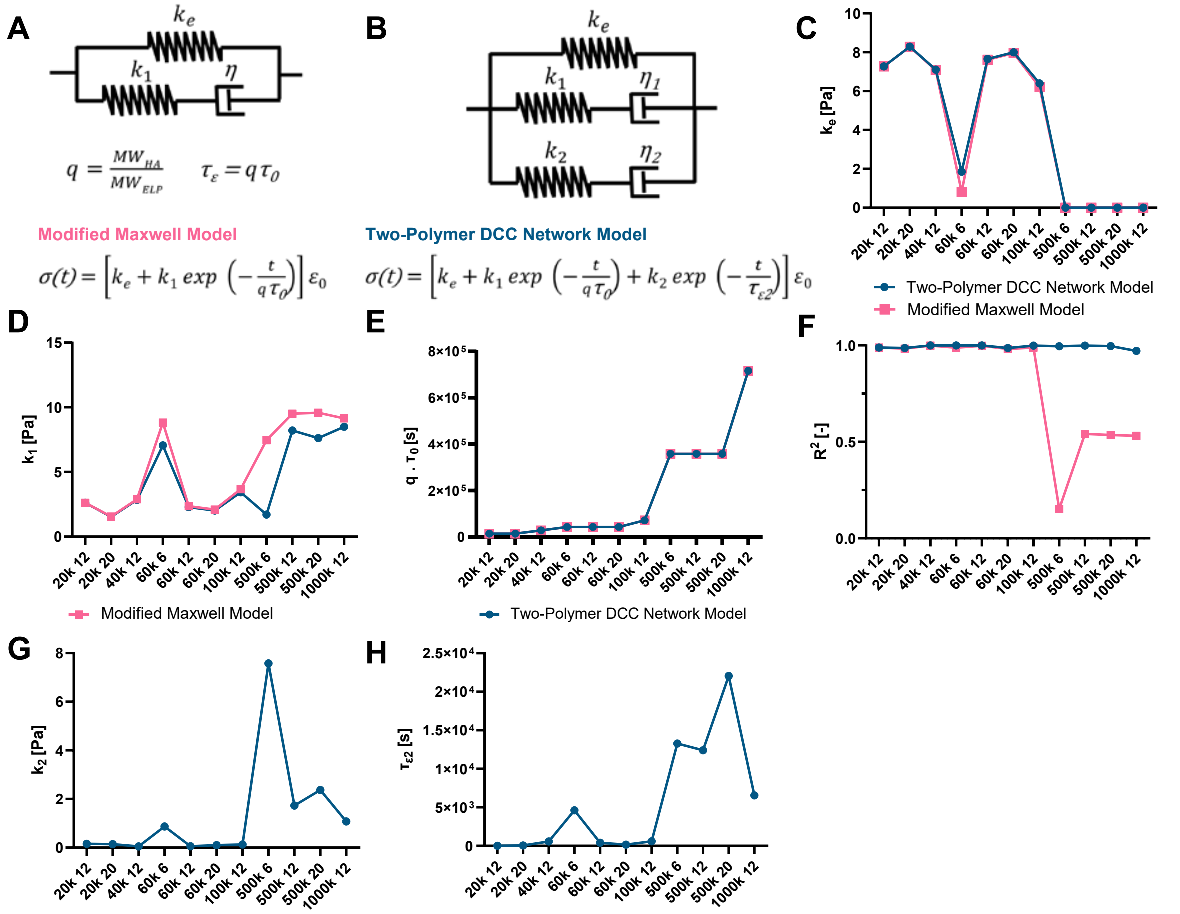


Figure S3: Comparison of fitted values for Two-Polymer DCC Network Model and Modified Maxwell Model

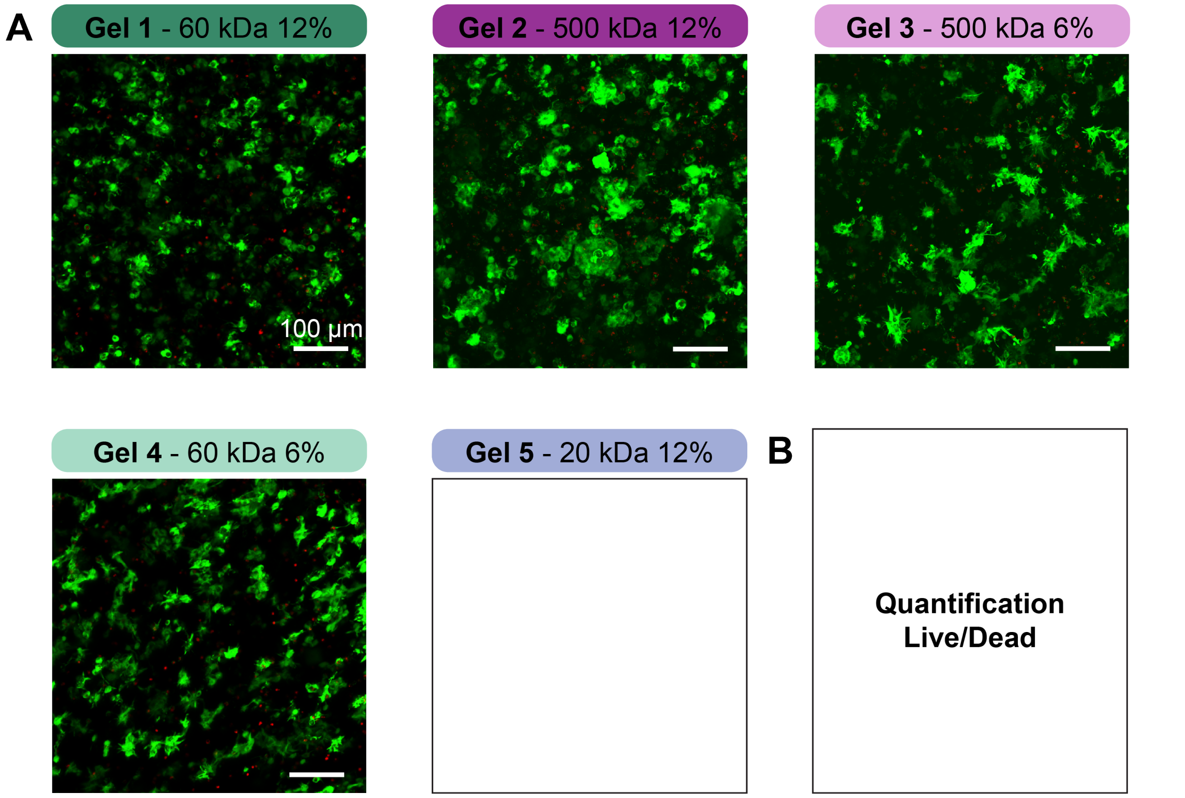


Figure S4: Live and Dead staining of hiPSC-derived NPCs within selected HELP formulations for 1 week
